# Mitigating Temozolomide Resistance in Glioblastoma via DNA Damage-Repair Inhibition

**DOI:** 10.1101/794115

**Authors:** Inmaculada C. Sorribes, Samuel K. Handelman, Harsh V. Jain

## Abstract

Glioblastomas are among the most lethal cancers, with a five year survival rate below 25%. Temozolomide is typically used in glioblastoma treatment; however, the enzymes APNG and MGMT efficiently mediate the repair of DNA damage caused by temozolomide, reducing treatment efficacy. Consequently, APNG and MGMT inhibition has been proposed as a way of overcoming chemotherapy resistance. Here, we develop a mechanistic mathematical model that explicitly incorporates the effect of chemotherapy on tumor cells, including the processes of DNA damage induction, cell arrest and DNA repair. Our model is carefully parameterized and validated, and then used to virtually recreate the response of heteroclonal glioblastoma to dual treatment with TMZ and inhibitors of APNG/MGMT. Using our mechanistic model, we identify four combination treatment strategies optimized by tumor cell phenotype, and isolate the strategy most likely to succeed in a pre-clinical and clinical setting. If confirmed in clinical trials, these strategies have the potential to offset chemotherapy resistance in glioblastoma patients, and improve overall survival.

## Introduction

Glioblastoma, or glioblastoma multiforme (GBM), is the most common and malignant of glial tumors, accounting for 60-75% of all astrocytomas. More than 90% of GBMs are estimated to develop over a period of only a few days or weeks, and are already grade IV cancers at the time of diagnosis [1]. The malignancy and lethality of GBM is driven by a rapid rate of cancer cell proliferation, coupled with a high degree of vascularity. Indeed, with improvements in leukemia survival since the 1980s, brain cancers are now the leading cause of childhood cancer death [2]. The first step in treating GBM is maximal resection or surgery, if possible. Radiation and chemotherapy with temozolomide (TMZ) may be administered subsequently [3, 4]. With standard treatment, the median survival of adults diagnosed with high-grade gliomas is less than 15 months, and fewer than 10% survive beyond 5 years; in children, GBM five year survival is below 25% [5]. Without treatment, GBM patients have a life expectancy of less than 3-4 months [3]. GBM-associated death rates remain high, in part, because the last few decades have produced modest advances in treatment. Consequently, the standard therapy for GBM remains palliative, rather than curative, and patients ultimately die from this disease [1].

Chemotherapy has proven to be effective against cancers in general; however, in the case of brain tumors, it has failed to produce sustained remission. GBM typically responds well in the first few cycles of TMZ administration. However, the emergence of resistance diminishes the cytotoxic effect of TMZ in subsequent cycles until there is little or no response to treatment [3]. Several factors may contribute to TMZ resistance (for a comprehensive review, see [6]). Here, we are concerned with resistance mediated by the efficient repair of treatment-induced DNA damage in cancer cells.

The chemical mechanism by which TMZ induces cell death is DNA methylation leading to double stranded breaks (DSBs) and thus to apoptosis. A range of mechanisms by which methylation causes DSBs are ably reviewed in [7]. The cell-killing potential of this methylation depends on the position within the purine bicyclic ring where the methyl-adduct is formed. The majority of TMZ-induced methylation sites are N7-meG (> 70%) and N3-meA (9.2%). Less frequently (< 6%) TMZ creates O6-meG adducts [8]. N7-meG and N3-meA contribute minimally to the cytotoxicity of TMZ as they are efficiently repaired via the base excision repair (BER) pathway, mediated by the alkylpurine-DNA-N-glycosylase (APNG) enzyme. O6-meG is more lethal for the cell, and is repaired via the methylguanine-DNA-methyltransferase (MGMT) pathway, mediated by the MGMT enzyme [8, 9]. Additionally, the mismatch repair (MMR) pathway may aid in DNA demethylation. However, the expression of MMR genes is linked to TMZ sensitivity [6]; therefore, this pathway is not considered here. The repair of TMZ-induced DNA damage reduces its therapeutic efficacy. Over-expression of both MGMT and APNG have been linked to resistance to alkylating agents, and are associated with lower survival rates in patients [10, 11], while MGMT methylation (a mark of reduced average expression) has been linked to improved survival. Consequently, it has been hypothesized that MGMT and APNG expression or methylation may be used as biomarkers for glioma response to treatment with alkylating agents such as TMZ [12, 13, 14].

Preventing the onset of chemotherapy-resistance by inhibiting MGMT or APNG in combination with TMZ administration represents an exciting new avenue of research with the potential for high impact in GBM treatment. Small molecule inhibitors of MGMT such as bortezomib, and APNG are in various stages of pre-clinical or clinical development, with a majority of this effort being directed at MGMT inhibition [7, 15]. With all of these new possibilities, a critical challenge in brain cancer therapeutics is the optimization of dosing and scheduling when alkylating agents such as TMZ are combined with DNA repair enzyme inhibitors. At the time of writing, no experimental studies could be found that consider the simultaneous inhibition of the two repair enzymes. Here, we propose a quantitative modeling framework that is used to conduct *in silico* pre-clinical and clinical trials with a view to predicting the potential of – and optimizing – such a combination.

Specifically, we develop a mathematical model of the response of GBM cells to TMZ administration, which incorporates TMZ-induced DNA damage, and its subsequent repair, at the level of intracellular molecular mechanisms. This allows us to simulate the effect of APNG and MGMT inhibition in a mechanistic, rather than phenomenological, manner. The model is extensively calibrated and validated versus available experimental data. We then optimize dosing and scheduling for the triple combination of TMZ and small molecule inhibitors of APNG and MGMT by simulating the treatment of phenotypically diverse GBM xenografts grown in virtual mice. Finally, the effect on patient survival of the combination with maximum anti-tumor potential is investigated by enrolling virtual human patients in an *in silico* clinical trial. Analysis of our results reveals critical features driving resistance to TMZ and evolution of tumor cell phenotypes under various treatment strategies. Our approach offers the key advantage of consistent comparison across strategies since a virtual patient may be treated with as many strategies as needed, allowing us to compare results without the confounding factor of inter-patient variability. Mathematical models have been employed extensively to describe the growth and treatment of gliomas (for instance, see [16, 17, 18, 19]; see [20] for a recent review). However, to the best of our knowledge, ours is the first to explicitly incorporate DNA methylation and repair pathways, and to consider DNA repair inhibition in combination with chemotherapy.

## Methods

### Mathematical Framework

Our modeling framework operates on multiple scales, from subcellular molecular interactions that govern outcomes at a cellular level, to the resultant population level behavior. A model schematic is shown in figure 1. Within the nucleus, TMZ induces methylation of nucleic acids, a process wherein a hydrogen atom on a DNA base is replaced by a methyl group (*R – CH*_3_). Methylation of DNA, in turn, leads to the formation of DNA adducts [8, 9]. Briefly, the active metabolite of TMZ binds to DNA reversibly, forming drug-DNA complexes with the rates of the forward reaction 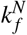 and 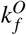 and reverse reaction 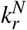 and 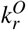 respectively. Subsequently, the methyl group on the drug is transferred irreversibly to DNA, resulting in adduct formation at rates 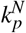 and 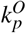, and TMZ is rendered inactive. Here, capital letters denote concentrations of intracellular species: *T*_*in*_, intracellular TMZ; *T*_*out*_ extracellular TMZ; *N*_7_, unmethylated DNA at N7-G or N3-A positions; *O*_6_, unmethylated DNA at O6-G positions; *D*_7_ and *D*_6_, position-specific drug-DNA complexes; *A*_7_ and *A*_6_, DNA adducts N7-meG/N3-meA and O6-meG, respectively; and *uT*_*in*_, inactive intracellular TMZ. We also explicitly include the transport of extracellular TMZ in and out of the cell at rates *γ*_*oi*_ and *γ*_*io*_, and the clearance of TMZ from the extracellular compartment at a rate *K*_*d*_. In our formulation, this process is approximated by the following reactions:

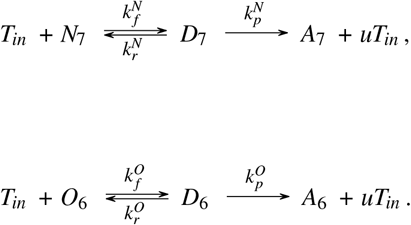

**Figure 1.**
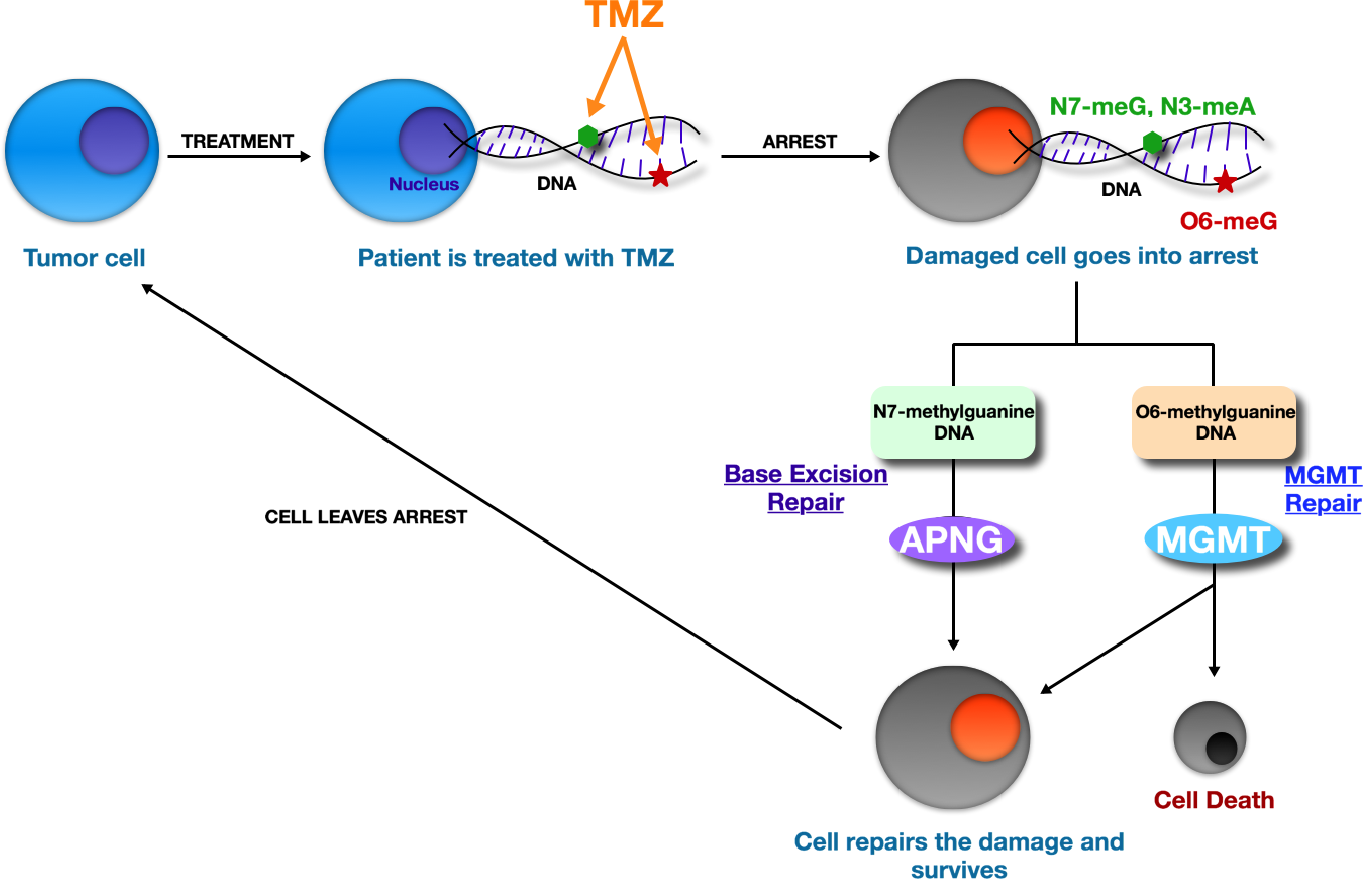
Model schematic. Treatment with TMZ causes DNA methylation leading to cell arrest. TMZ-induced methylation sites are the N7-meG and N3-meA (repaired by APNG-mediated BER pathway), and O6-meG (repaired by MGMT pathway). If DNA damage repair is successful, the cell recovers to the proliferating pool, or undergoes apoptosis otherwise.

Assuming this, the above chemical reactions can be translated into mathematical equations assuming mass action kinetics:

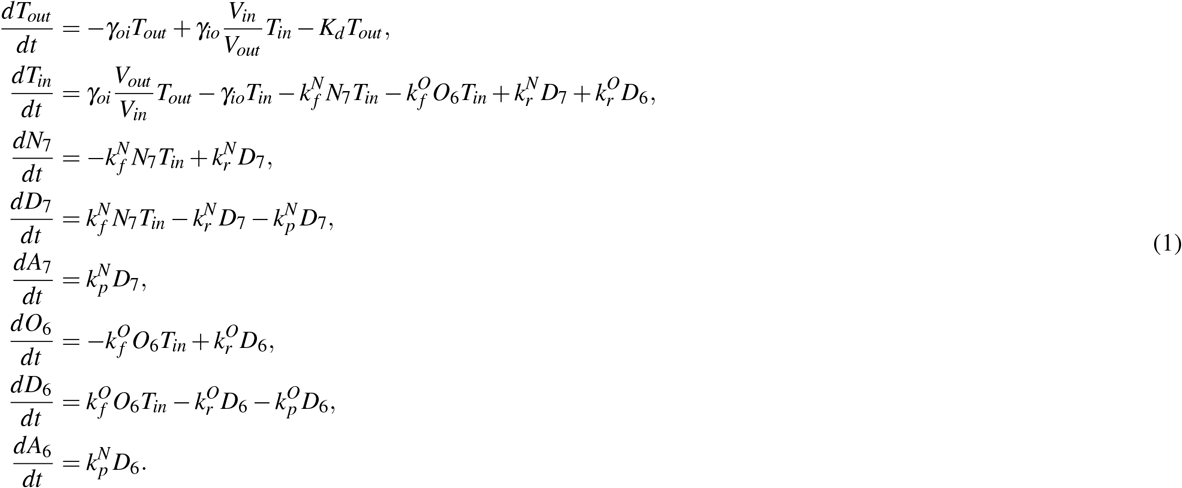

Once DNA damage exceeds a critical threshold, the cell enters a state of arrest and different DNA repair mechanisms are triggered depending on the site of damage. For clarity, subcelullar species in arrested cells are represented with *. We focus on the BER and the MGMT pathways. The BER enzyme APNG is responsible for repairing N3-meG and N7-meA DNA adducts in arrested cells. APNG binds reversibly to the methyl group, with the rate of the forward reaction 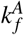 and reverse reaction 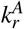 creating a complex. The damage is successfully repaired at a rate 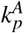, resulting in demethylated DNA in the arrested cell. The following reaction represents our assumptions

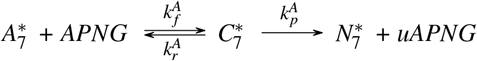

On the other hand, when MGMT encounters an O6-meG adduct in an arrested cell, MGMT transfers the alkyl group to an internal cytosine residue at a rate 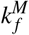, acting as a transferase and the receptor of the alkyl group. We consider that the transfer of the methyl group can be unsuccessful, at a rate 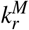. After successfully receiving the alkyl group from the guanine MGMT inactivates itself becoming a suicidal protein [21]. When the methyl group from the O6 position of the guanine is removed, at a rate 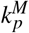, the guanine is restored to its standard form without breaking DNA strands. The reactions read as follows

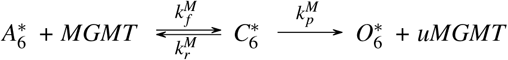

Assuming mass action kinetics, and including production and degradation of species when appropiate, the avobe reactions lead to the equations governing DNA repair:

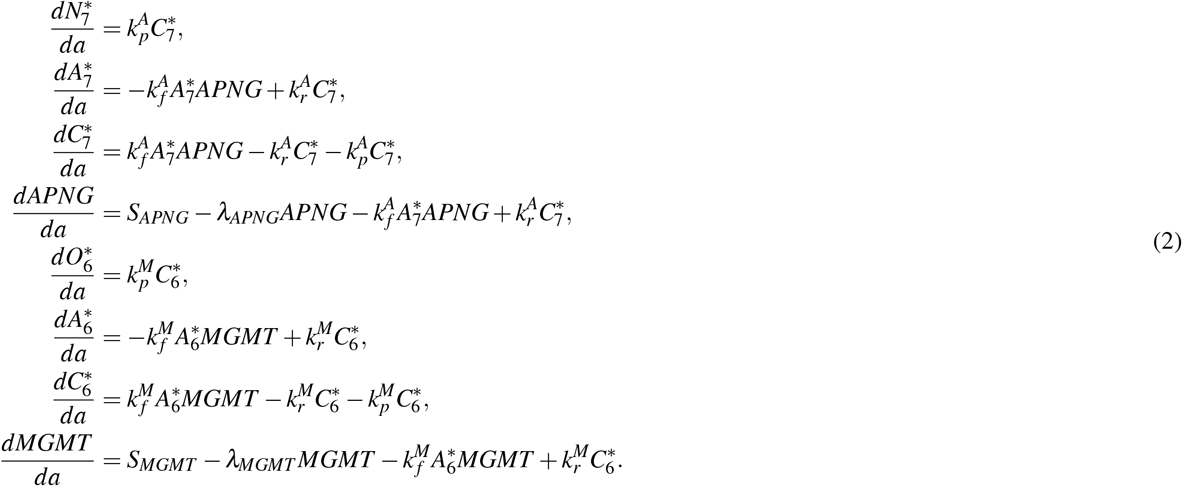

Here: *a* is a time variable that represents how long a cell has spent in the arrested compartment; 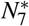, represents the concentration of demethylated DNA at a N7-G position; 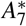, represents the concentration of DNA adducts N7-meG; 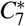, represents the concentration of the complex formed by APNG binding to N7-meG; *APNG* represents the concentration of APNG; 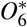, represents the concentration of demethylated DNA at an O6-G position; 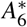, represents the concentration of DNA adducts O6-meG; 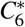, represents the concentration of the complex formed by MGMT binding to O6-meG; and *MGMT* represents the concentration of MGMT. Further, *S*_*APNG*_ represents the production of APNG; *S*_*MGMT*_, represents the production of MGMT; *λ*_*APNG*_, represents the degradation rate of APNG; and *λ*_*MGMT*_ represents the degradation rate of MGMT. In reality these pathways are very complex. In order to avoid an over-parameterized model, or lose focus on the essential aspects, we considered a simplification of the actual pathways.

At the population level, the tumor is compartmentalized into proliferating and arrested cells. Following experimentally observed growth patterns, proliferating cells are assumed to grow either logistically, when simulating *in vitro* cell growth inhibition assays and *in vivo* xenograft assays, or exponentially, when simulating GBM growing *in situ*. TMZ application induces DNA damage, leading to cell cycle arrest and triggering of the BER and MGMT pathways [6]. An age-structure is imposed on the arrested compartment to account for the time taken from induction of DNA damage to cell death, or recovery following successful completion of damage repair. Noting that TMZ specifically targets actively proliferating cells, inducing arrest in G2/M phase [9, 22], cells in the arrested compartment are assumed to be unaffected by its further application.

Mathematically, *P*(*t*) represents the number of proliferating tumor cells at time *t*, and *A*(*t, a*), the number of arrested cells at time *t* that have been arrested for *a* units of time. The following equations represent the population-level processes described above:

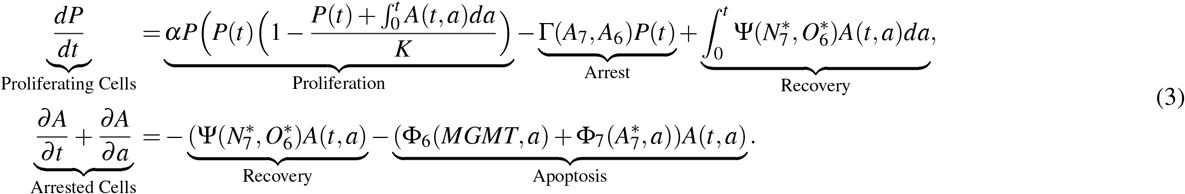

Where *α* represents the proliferation rate of tumor cells; *K*, represents the carrying capacity (when needed); Γ, represents the rate of cell arrest; Ψ, represents the rate of cell repair; Φ_6_ represents the rate of cell death corresponding O6-meG; and Φ_7_ represents the rate of cell death corresponding to N7-meG. We remark that when a cell is arrested, TMZ does not further cause DNA damage to it, and DNA repair only occurs in the arrested compartment.

The subcellular and population scales are connected by decisions made at the cellular level, represented in the above equations with capital Greek letters. For instance, cells enter a state of arrest at a rate Γ, which is assumed to depend on the level of DNA damage. From an arrested state, cells may recover to the proliferating pool at a rate Ψ, if DNA damage repair has been successful, which is assumed to be proportional to the fraction of DNA that has been repaired. The constants of proportionality are *μ* and *ρ* respectively. Arrested cells may also undergo apoptosis at rates Φ_6_ and Φ_7_, taken to be functions of the levels of damaged DNA and repair enzymes, and the amount of time spent in an arrested state. It has been shown that enhanced cytotoxicity is associated with increased time of arrest [23, 24, 25, 26, 27]. Thus, we assume that the rate of cell death caused by both N7-meG and O6-meG is comprised of a smooth step function that considers cell death as a function of time of arrest, where *c*_1_ is the rate of cell death caused by a long arrest time, and where *b*_1_ and *b*_2_ are the rates of cell death caused by DNA damage. We denote the characteristic waiting time of arrested cells until apoptosis starts to occur by *a*_0_. Further, it has been shown that cells that express MGMT are protected against apoptosis after treatment with TMZ [28], thus we consider the rate of death caused by O6-meG to be a function of *a* and the concentration of MGMT. On the other hand, when N3-meA and N7-meG are present in high levels they trigger apoptosis [28], thus we choose the rate of death Φ_7_ to be a function of *a* and concentration of N7-meG and N3-meA. In particular,

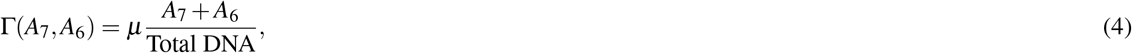

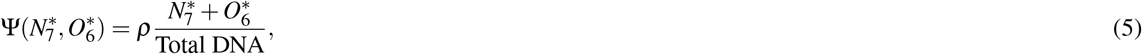

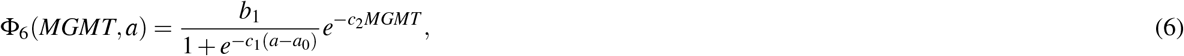

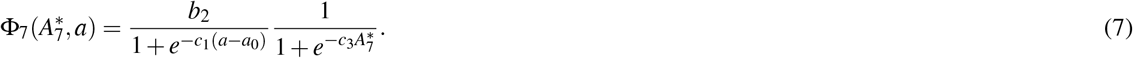

The various model variables and parameters are listed with units and biological explanations in the Supplemental Information, tables S1 and S2.

These equations are closed by imposing initial and boundary conditions. Here we present the most general initial conditions that close the system. We note that initial conditions may vary depending on the experimental settings. For instance, experiments conducted *in vitro* or in xenograft will assume an initial population of tumor cells of 10^5^, which is consistent with current clinical practice [29, 30, 31, 32, 33]. However, for simulations of GBMs growing in humans we assume an initial tumor measuring 1 cm [34] (corresponding to 5 × 10^8^ cells considering that GBM cells measure 12 – 14*μ*m [35]). In general *t* = 0 will represent the first time TMZ is administered and this will give the following initial and boundary conditions of equations (3)

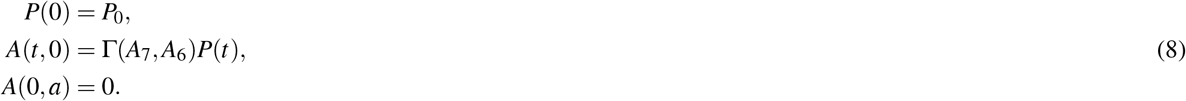

The initial conditions in the DNA damage equations (1) are

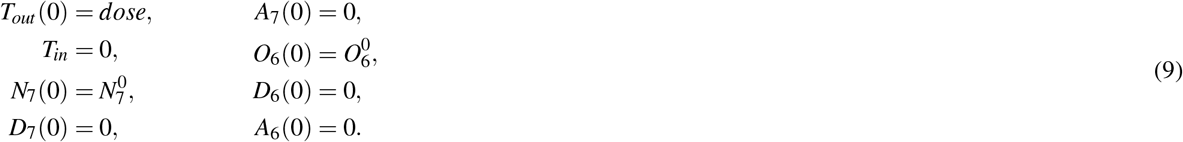

Once cells become arrested the initial conditions for equations (2) are

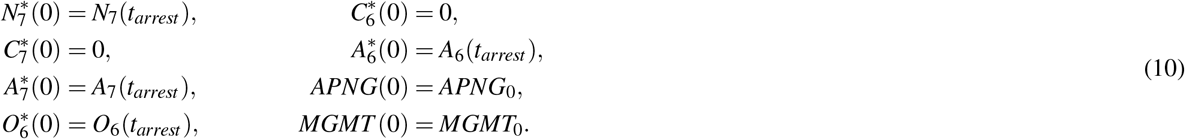

The initial value of *APNG* and *MGMT* will depend on the specific cell type, and they are estimated later on. If a tumor cell returns to the proliferating compartment we assume that all DNA has been repaired and *N*_7_ and *O*_6_ return to their initial value, while *A*_7_, *A*_6_ are reset to be zero.

### Structural identifiability and estimation of model parameters

Our model has 31 parameters that need to be estimated from available experimental data. With such a large number of unknowns, it is crucial to check whether the estimation problem is well-posed [36]. This question is addressed by conducting a structural identifiability analysis, a process which determines whether indistinguishable outputs of the model imply uniqueness of parameters, disregarding issues related to data quantity or quality [37]. This analysis reveals the maximum possible information about parameters a given type of data contains and is therefore essential if we are to have any degree of confidence in parameter estimates and, by extension, model predictions.

Here we use the Matlab toolbox GenSSI 2.0, which uses a series approach to determine structural identifiability and presents the results using identifiability tableaus (for details, see [38]). Briefly, Lie derivatives are employed to arrive at a system of equations for model parameters in terms of model variables. The analysis has three possible outcomes for each parameter. A given parameter is deemed globally identifiable if the resulting system can be solved uniquely for it, that is, the parameter can be uniquely determined from the system output. A parameter is locally identifiable if the above result holds in a neighborhood of it. Finally, a parameter is unidentifiable if infinitely many values of it yield the same model output [37]. All parameters were found to be globally or locally structurally identifiable, thus validating the structure of our proposed model.

We next extensively calibrated parameters using available clinical data from a wide array of experiments. Moreover, model output was found to match experimental data that were excluded from the fitting process, thus validating our model formulation. A detailed description of the structural identifiability analysis, clinical data, parameter estimates, and model validation is provided in the Supplemental Information. Identifiability tableaus for this analysis are presented in figure S1 F-H, best fits are shown in figure S2, model validation in figure S3, with the parameter value estimates recorded in tables S3 and S4.

### Sensitivity analysis and energy constraints yield phenotypically diverse virtual GBM cell lines

Predicting optimal treatment protocols not only requires a validated mechanistic model, but also a phenotypically diverse population of cells that is representative of different responses to treatment. We generate such a cohort of virtual cell lines by first identifying, by means of local and global sensitivity analyses, which features – or model parameters – are critical drivers of resistance to TMZ. Next, diverse cell phenotypes are generated by allowing these features to vary randomly, keeping in mind that all cellular processes require energy, only a limited amount of which is available to each cell.

Significant variance has been observed across cell populations in proliferation rates (or doubling times) [39], and repair enzyme expression levels [40]. Further, repair enzyme turnover time-scales may affect how fast DNA damage repair is completed. Therefore, in our sensitivity analysis, we focus on the rates of: cell proliferation (*α*); repair enzyme expression (*S*_*MGMT*_, *S*_*APNG*_); and repair enzyme degradation (*λ*_*APNG*_, *λ*_*MGMT*_).

We first conducted a local sensitivity analysis, the results of which are presented in figures 2 and S4. Following the experimental protocol in [41], cell growth inhibition assays were simulated, with 10^5^ cells cultured in the presence of 500*μ*M TMZ for 1 week, and surviving cells counted at the end of the experiment. Figure 2A reveals that cell survival increases as repair enzyme expression rates are increased, this effect being more pronounced as MGMT expression is varied for a fixed level of APNG expression. Interestingly, cells expressing high levels of MGMT show similarly high survival rates even if APNG expression is suppressed. We next investigate the effect of varying the rates of cell proliferation and repair enzyme turnover on cell survival. For each parameter, four distinct cell phenotypes are considered, based on whether or not MGMT and APNG are expressed. From figure 2B, we observe that in cells expressing at least one repair enzyme, increasing the rate of cell proliferation – or, equivalently, decreasing cell doubling time – is associated with improved cell survival, and hence greater resistance to TMZ. Further, as can be seen in figure 2C, whenever cells express MGMT, cell survival increases as the rate of MGMT turnover increases. On the other hand, figure 2D reveals that the rate of APNG turnover has little impact on cell survival. This is because we account for the experimentally observed [28] pro-survival effect of MGMT on TMZ treated cells. In all cases, cells not expressing either repair enzyme are most sensitive to TMZ and their survival is minimally affected by varying any of the considered parameters.

**Figure 2.**
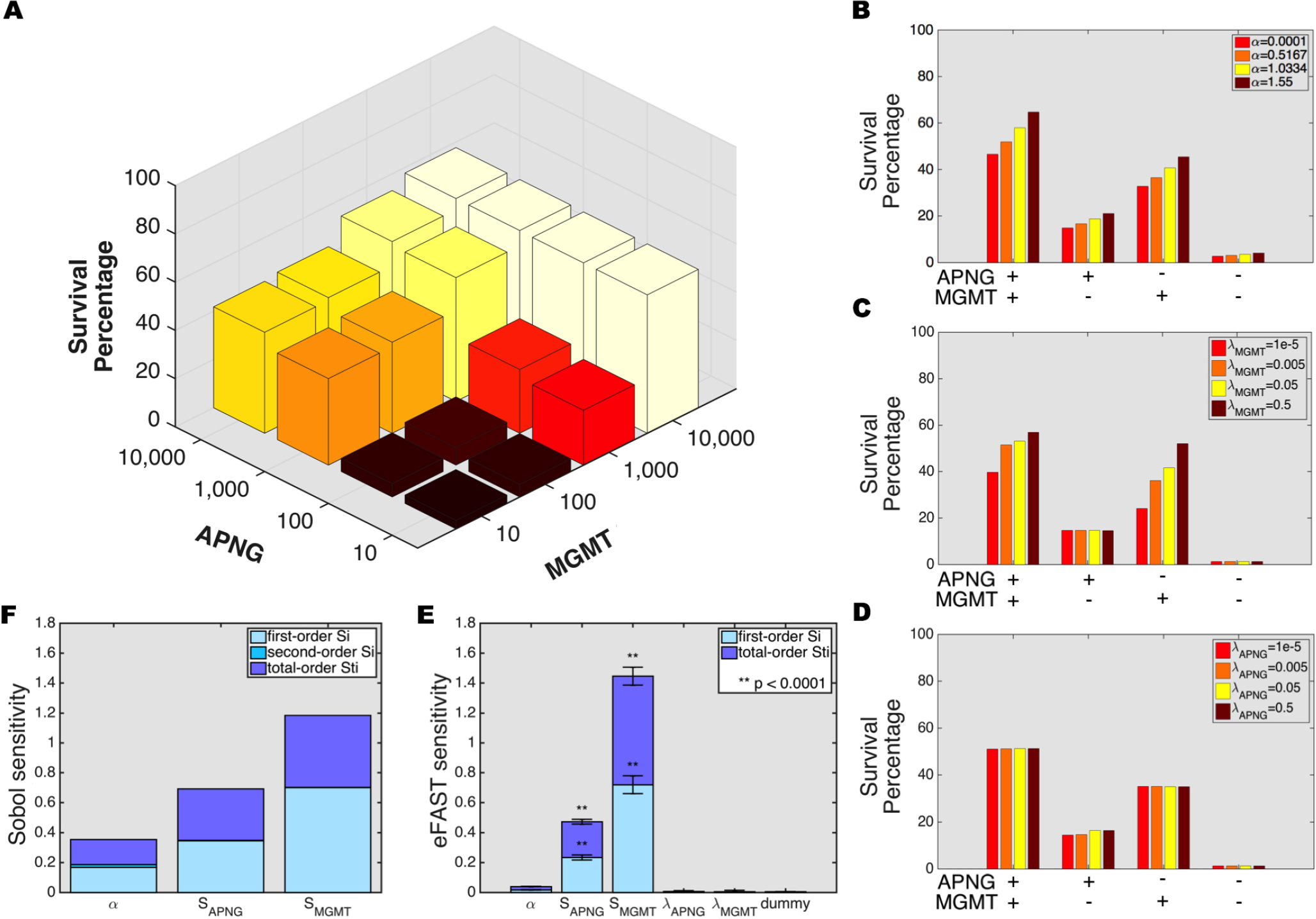
Local and global sensitivity analyses. A, 3D histogram of survival percentage of cells post-TMZ treatment in a cell growth inhibition assay as the expression levels of APNG (*y*-axis) and MGMT (*x*-axis) are varied. B, survival percentage of cells as proliferation rate is varied in cell lines with MGMT and/or APNG expression turned on/off. C,D, survival percentage of cells as MGMT turnover rate (C) or APNG turnover rate (D) is varied in cell lines with MGMT and/or APNG expression turned on/off. E, eFAST sensitivity indices of various model parameters (*α*, proliferation rate, *S*_*APNG*_ and *S*_*MGMT*_, rates of APNG and MGMT expression, respectively and *λ*_*APNG*_ and *λ*_*MGMT*_, rates of APNG and MGMT degradation, respectively). A dummy variable is used to perform a t-test. F, Sobol sensitivity indices of the parameters *α*, *S*_*APNG*_, and *S*_*MGMT*_. See also figure S4.

Local sensitivity analysis ignores the effect of any correlation between various parameters, and hence, the biological processes they represent. Therefore we also performed a global sensitivity analysis with all parameters being varied simultaneously, using the eFAST and Sobol methods [42]. Figures 2E and 2F plot eFAST first- and total-order indices and Sobol’s first-order and total effect indices, respectively, for the various parameters considered. We see that resistance to TMZ is maximally affected by MGMT expression, followed by APNG expression and, to a lesser extent, cell proliferation rate. The rates of turnover of *both* enzymes have a minimal impact on cell survival, with their sensitivity indices indistinguishable from that of a dummy variable (figure 2E).

From the above analyses, we conclude that the rates of MGMT and APNG expression and cell proliferation may be significant determinants of tumor cell sensitivity to TMZ. Therefore, virtual cell lines with phenotypes capable of capturing diverse responses to TMZ are created by allowing *S*_*MGMT*_, *S*_*APNG*_ and *α* to vary randomly, keeping all other model parameters fixed. However, cellular activities such as proliferation and repairing DNA damage require energy, and tumor cells have to balance the allocation of limited resources towards such activities. Motivated by the approach of Nagy et al. [43], we derive an energy constraint inequality to which all cells must adhere. This places bounds on how *S*_*MGMT*_, *S*_*APNG*_ and *α* may vary relative to each other.

Specifically, tumor cells perform the following activities: cell division; APNG and/or MGMT expression; and maintenance of normal physiological function (inclusive of all other cellular processes). Each activity requires the production of relevant cellular material, for which energy – provided by ATP – is consumed. Based on data relating to ATP reserves maintained in a cell, and typical rates of glycolysis and ATP hydrolysis, an upper bound for the energy available to a cell (*M*_*E*_*C*__) is derived, resulting in the following energy constraint inequality. We define energy as the effort required to create 1*μ*M material in 1 minute, where effort relates to the rate of ATP consumption (see Supplemental Information for details):

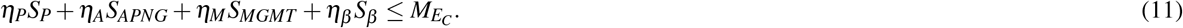

Here: *η*_*P*_, *η*_*A*_, *η*_*M*_ and *η*_*β*_ are efforts related to proliferation, APNG production, MGMT production and maintenance, respectively. *S*_*P*_, taken to be proportional to *α*, and *S*_*β*_ are the production rates of cellular material needed for division and maintenance, respectively. Virtual cell lines are generated by allowing *S*_*P*_, *S*_*APNG*_ and *S*_*MGMT*_ to vary, whilst ensuring the above equation is satisfied. The maximum threshold allowed in virtual cells is based of T98G cells, an established GBM cell line known for its virulence. We remark that absent such a constraint, cells with the least doubling time and maximum repair enzyme expression would dominate. However, such cells would consume biologically unrealistic amounts of energy.

## Results

### MGMT and APNG expression, rather than cell doubling time, are indicators of resistance to TMZ

Next, we simulate xenograft treatment assays to determine the relative contribution to TMZ resistance of the key features identified above. Briefly, 500 ‘virtual’ mice received intracranial implantations of 10^5^ GBM cells. Each such implantation comprised equal numbers of 10 randomly chosen cell lines, thereby ensuring phenotypic heterogeneity in the resultant xenografts. At day 0, each animal was treated with a single dose of 500*μ*M TMZ, and tumor volume and composition recorded 1 week later. Cell phenotypes were classified by the percentage of the tumor they occupied at the end of the experiment with those cell lines assigned to quartile 1 that occupy between 0-25% of the tumor, those to quartile 2 that occupy between 25-50% of the tumor, and so on. Thus, cell lines belonging to quartile 4 for instance, dominate the tumor and represent phenotypes most resistant to TMZ.

The resultant tumor composition profiles are shown in figure 3A-C as RDI (Raw data, Descriptive statistics, and Inferential statistics) plots. Figure 3A reveals that faster proliferating cells are more resistant to TMZ, with the smoothed density of cells in quartile 4 showing a top-heavy distribution. However, the differences in the central tendencies of cell proliferation rates across the various quartiles are minimal. Indeed, for cell lines occupying quartiles 1-3, these differences are not statistically significant. Therefore, we conclude that cell doubling time alone cannot guarantee resistance to TMZ.

**Figure 3.**
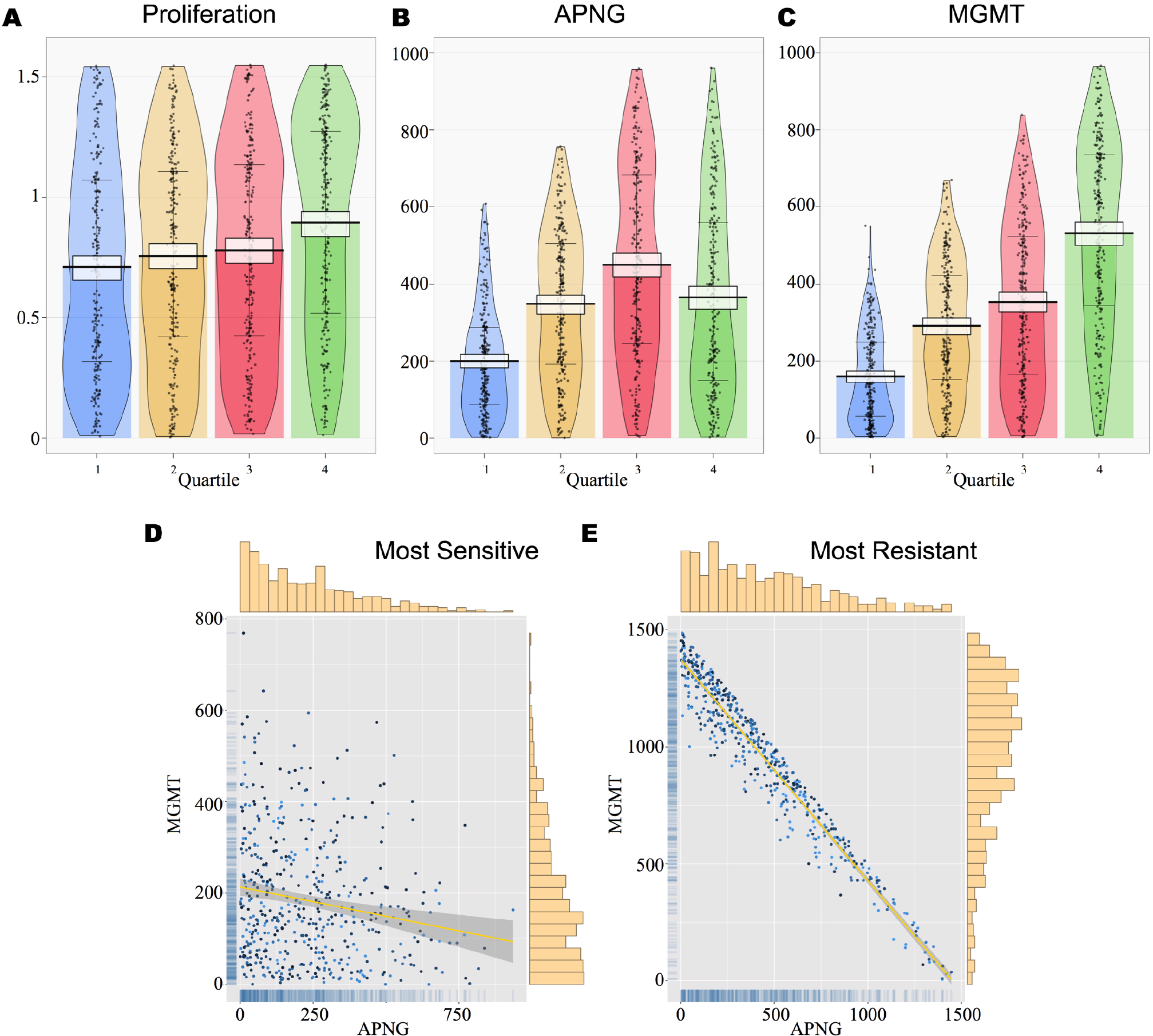
Heteroclonal tumor xenograft response to a single dose of TMZ. A-C, RDI (Raw data, Descriptive statistics, and Inferential statistics) plots showing raw data points (black dots), central tendencies (vertical bar), smoothed densities (irregular colored regions), and Bayesian highest density intervals (white rectangles). Tumor cell lines are divided into quartiles according to the percentage of the tumor they occupy at the end of the experiment: quartile 1, 0-25%; quartile 2, 25-50%; quartile 3, 50-75%; and quartile 4, 75-100%. Cell lines in quartile 1 are most sensitive to TMZ, while those in quartile 4 are most resistant. Three cellular phenotypes are considered: A, rate of cell proliferation; B, APNG expression; and C, MGMT expression. D,E, scatter plots of APNG expression versus MGMT expression in the most TMZ-sensitive cells (D) and most TMZ-resistant cells (E). The color of the data dots is a gradient based on proliferation rate with darker dots representing cells with smaller doubling time. Linear correlation is shown as yellow lines, and corresponding 95% confidence intervals shown as shaded areas. Distributions of enzyme expression are shown as marginal histograms (top, APNG and right, MGMT) and marginal rugs (left, MGMT and bottom, APNG).

The expression levels of repair enzymes reveal more differentiating attributes (figure 3B,C). As expected, cell lines most sensitive to TMZ (quartile 1) express the least amounts of both APNG and MGMT, with clear bottom-heavy distributions and narrow 95% highest density intervals (HDIs) around the central tendency. Interestingly, figure 3B reveals that although the central tendency of APNG expression increases from quartile 1 to 3, the most resistant cell lines (quartile 4) show a significant decrease in the most credible values of APNG expression. The reason for such an unexpected decline becomes clear when we look at the expression of MGMT in figure 3C. We observe that resistance to TMZ increases with MGMT expression monotonically. Further, the smoothed density of cells in quartile 4 is top heavy, indicating that the most resistant cell lines invest maximally in MGMT expression. This comes at the cost of APNG expression due to the imposed energy constraint. Thus we conclude that MGMT– and, perhaps to a lesser extent, APNG–expression are reliable predictors of TMZ-resistance.

Further verification of our conclusion comes from looking at scatter plots of APNG versus MGMT expression in the most sensitive (quartile 1) and most resistant (quartile 4) cell lines, shown in figure 3D-E, respectively. Linear regressions, shown in yellow, reveal that rates of APNG and MGMT expression correlate poorly in TMZ-sensitive cell lines (adjusted R-squared: 0.03) while resistant cell lines maximize their efforts towards repairing cell damage. In these cells, a clear linear trend in APNG versus MGMT expression is observed (adjusted R-squared: 0.94), with high values of MGMT expression favored. Dots represent individual cell lines, with darker colors corresponding to cells with higher proliferation rates, and vice versa. Neither plot shows a discernible pattern in the color distribution of dots, indicating a lack of correlation between resistance to TMZ and cell proliferation rate.

### Optimizing TMZ administration with APNG and MGMT inhibition

We now predict optimal drug dosing and scheduling protocols when small molecule inhibitors of APNG and MGMT are co-administered with TMZ. Briefly, 500 phenotypically diverse GBM cell lines were generated and cell growth inhibition assays initiated as described previously. A single cycle of treatment was simulated, with drug administration possible on days 1-5, followed by a rest period from days 6-28. This is consistent with typical clinical protocols when administering TMZ. A genetic algorithm [44] was employed to arrive at optimal treatment strategies specific to individual cell lines, with variability allowed in the daily TMZ dosage, and a decision made whether or not to co-administer APNG and/or MGMT inhibitors. The algorithm ranked treatment strategies based on their reduction of tumor burden at the end of the protocol. Further details of this process are provided in the Supplemental Information.

An analysis of the optimal protocols thus obtained reveals a natural grouping which is based on tumor cell phenotype. Specifically, we can classify the cell lines into 4 cohorts distinguished by the expression of each enzyme. For instance, cells expressing high levels of both repair enzymes are assigned to the ‘APNG+/MGMT+’ cohort, and so on. The “optimal” protocol for each cohort is then obtained by averaging the TMZ dosage across cell lines in that cohort, and finding the mode of the decision whether or not to inhibit APNG and/or MGMT. The resultant protocols are summarized in table 1.

**Table 1.** Description of treatment strategies. Standard, and optimal treatment strategies identified by GBM cell phenotype. Note that the specific daily dose of TMZ varies between strategies. Here, SMI stands for small molecule inhibitor.

|  |  | Treatment Strategy | TMZ | SMI APNG | SMI MGMT |
| --- | --- | --- | --- | --- | --- |
|  |  | Standard TMZ | Constant dose days 1 – 5<br>Rest days 6 – 28 | None | None |
|  |  | Optimal TMZ | High dose days 1, 3, and 5<br>Low dose days 2, and 4<br>Rest days 6 – 28 | None | None |
| Cell Phenotype | APNG–/MGMT– | A | High dose days 1, 3, and 5<br>Low dose days 2, and 4<br>Rest days 6 – 28 | Days 1 – 5 | Day 1 and 5 |
|  | APNG+/MGMT– | B | High dose days 1, 3, and 5<br>Low dose days 2, and 4<br>Rest days 6 – 28 | Days 1 – 5 | Days 2, 3, and 4 |
|  | APNG–/MGMT+ | C | High dose days 1, 3, and 5<br>Low dose days 2, and 4<br>Rest days 6 – 28 | Days 1 – 5 | Days 1, 2, 3, and 4 |
|  | APNG+/MGMT+ | D | High dose days 1, 3, and 5<br>Low dose days 2, and 4<br>Rest days 6 – 28 | Days 1 – 5 | Days 1 – 5 |

In all cases, alternating the daily dose of TMZ, with a high dose followed by a low dose, is predicted to be optimal. This is in contrast with typical clinical practice where a fixed dose is administered daily, referred to here as standard TMZ. Indeed, the experiments of Wick et al. [45] provide indirect validation for this prediction, where it was observed that cells over-expressing MGMT were more susceptible to an alternating TMZ dose.

Interestingly, optimal protocols in every case require APNG inhibition to be applied on *all* treatment days. This is surprising since N7-meG and N3-meA typically contribute minimally to the cytotoxicity of TMZ [8]. However, our model accounts for the fact that high levels of N7-meG adducts trigger apoptosis [28]. Further, APNG is predicted to be highly expressed in cell lines with a greater than average degree of TMZ resistance (see figure 3B, quartile 3). Together, these features explain the predicted optimal schedule of APNG inhibitors. In fact, it is the scheduling of MGMT inhibitors that distinguishes the four cohorts, with cell lines in the APNG+/MGMT+ cohort – and hence, most resistant to TMZ – requiring MGMT inhibition on all treatment days. It is worth noting that the optimal scheduling of MGMT inhibition is not dictated exclusively by its expression (compare rows 3-4 or 5-6 in table 1), but seems to also depend on APNG expression. This highlights a key advantage of our approach, namely, the mechanistic model of subcellular response to TMZ that underpins our simulations captures unexpected and potentially non-linear interdependencies between the two DNA damage repair pathways.

A comparison of how cell lines in each cohort respond to standard and optimal TMZ monotherapy, and optimal combination therapy, is shown in figure 4. For each cohort and for each treatment strategy, surviving tumor cell number together with 95% confidence intervals are plotted versus time. In general, optimal TMZ performs marginally better than standard TMZ. Maximum cell growth inhibition, driven by a high degree of apoptosis early on, is achieved when TMZ is combined with APNG and MGMT inhibitors. Cell survival 7 days post-treatment initiation is comparable across strategies when cell lines express low levels of MGMT (figure 4A,B). However, combination therapy is most effective when treating cells expressing high levels of MGMT (figure 4C,D).

**Figure 4.**
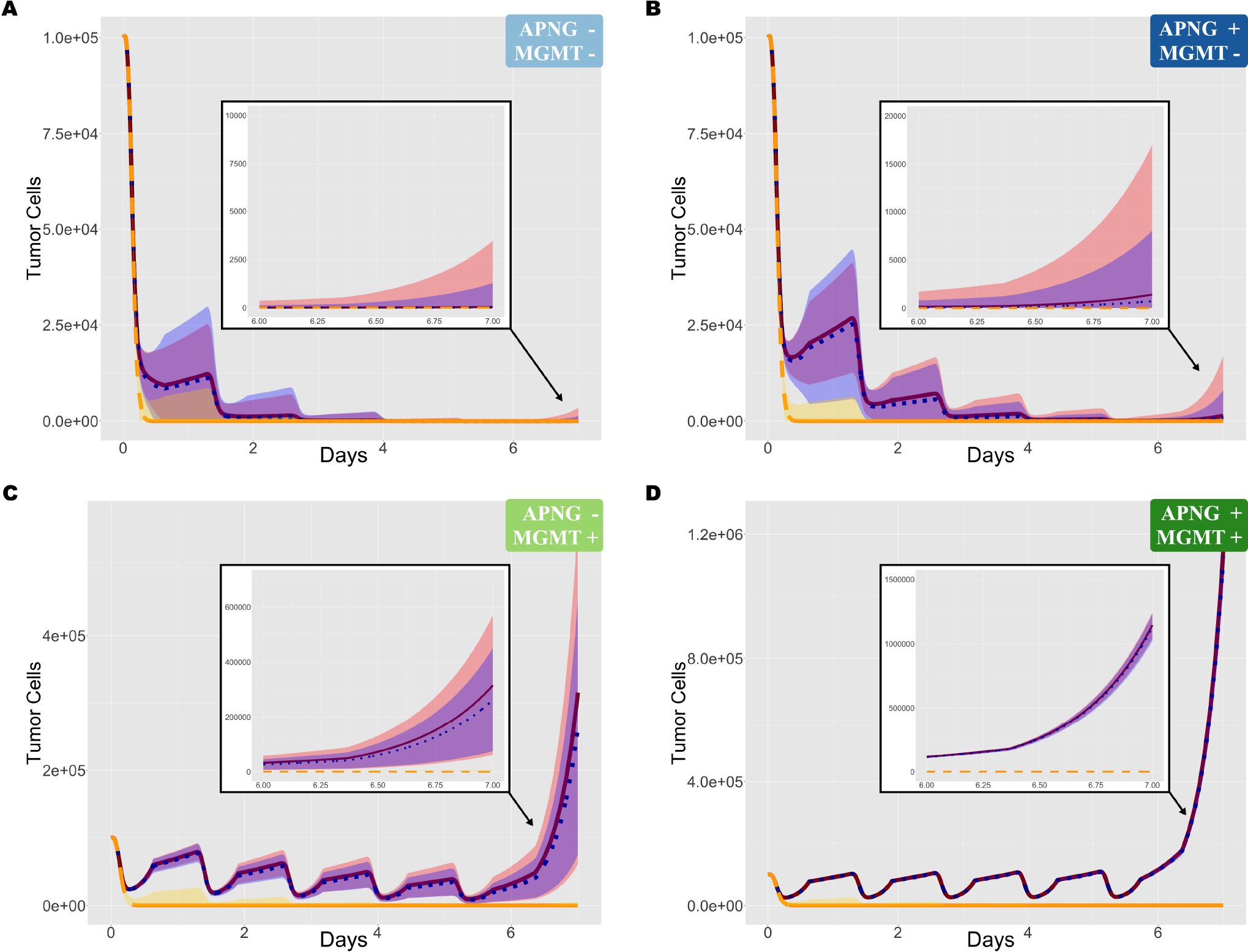
Cell growth inhibition in response to various treatment strategies. Mean values and 95% confidence intervals of surviving cell numbers when cell growth inhibition assays are simulated with: standard TMZ (solid red line, red shaded area); optimal TMZ (dotted blue line, blue shaded area); and optimal combination treatment (dashed yellow line, yellow shaded area). A, cells expressing low levels of both repair enzymes (APNG−/MGMT−). B, cells expressing high levels of APNG and low levels of MGMT (APNG+/MGMT−). C, cells expressing low levels of APNG and high levels of MGMT (APNG−/MGMT+). D, cells expressing high levels of both repair enzymes (APNG+/MGMT+).

### Treatment strategy D outperforms all others in a virtual pre-clinical trial

When treating GBM *in situ*, the expression levels of MGMT and APNG may not be known *a priori*. Indeed, these may vary significantly even within a tumor. Therefore, we identify the protocol with maximum inhibitory potential when treating tumors comprising a heterogeneous population of cells. For this, a pre-clinical trial was conducted *in silico* as follows. 500 virtual mice, with heteroclonal tumor xenografts established as described previously, were randomly assigned to one of the optimal treatment strategies (A, B, C, or D listed in table 1). Each mouse received a single cycle of treatment, and tumor volumes were recorded periodically. The resulting averages of the fold-change in tumor volume relative to pre-treatment, together with 95% confidence intervals, are shown as a waterfall plot in figure 5.

**Figure 5.**
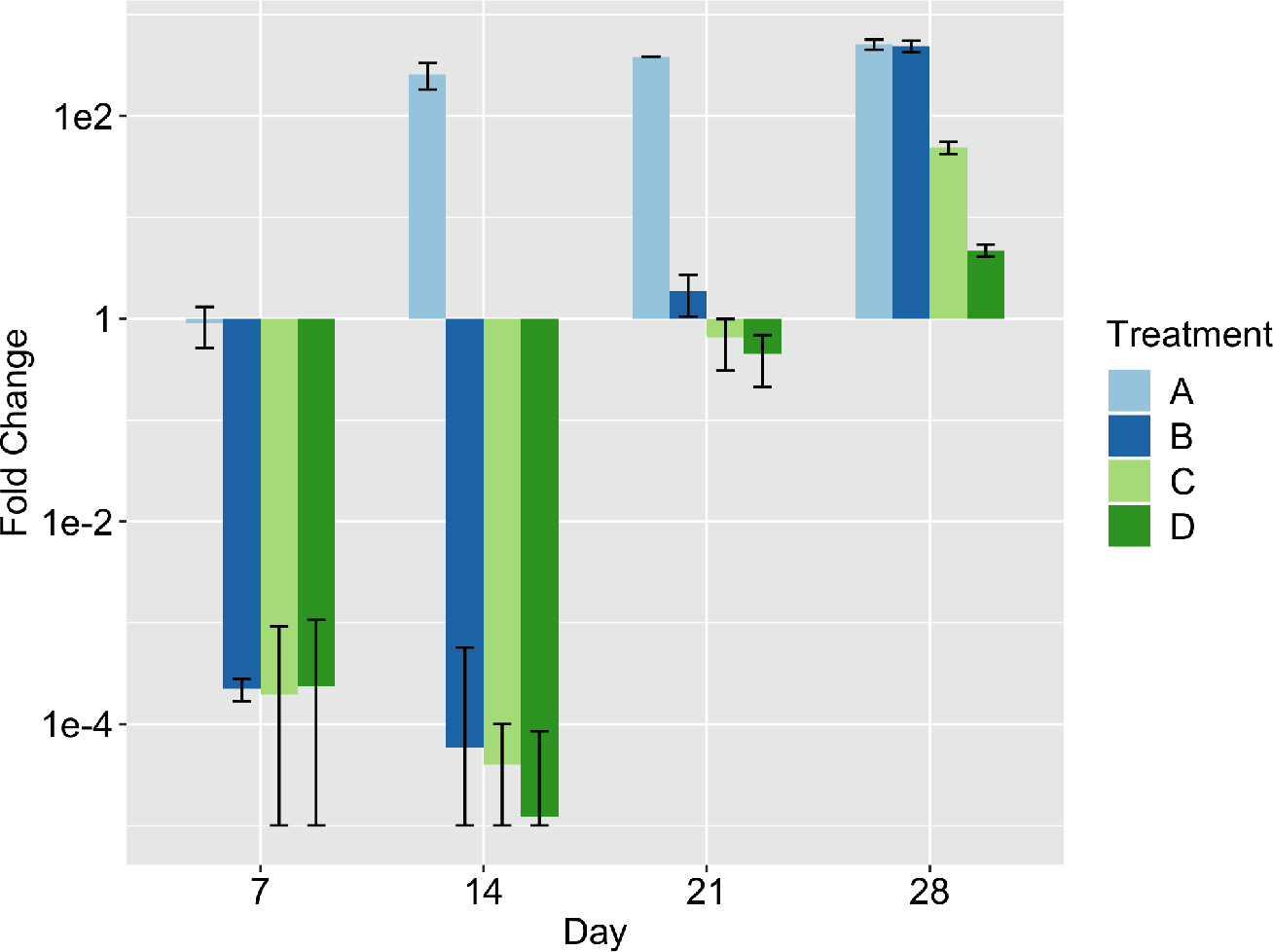
Heteroclonal xenograft response to optimal treatment protocols identified in table 1. Mice with heteroclonal tumor xenografts were randomly assigned to one of the strategies A, B, C, or D. The mean values of fold-change in tumor volume relative to pre-treatment are shown at days 7, 14, 21, and 28 post-treatment initiation. Error bars represent 95% confidence intervals.

Even at week 1, strategy A, optimal for APNG /MGMT cells, has failed to induce a significant reduction in tumor volume. In contrast, strategies B, C and D, optimal for APNG+/MGMT, APNG /MGMT+ and APNG+/MGMT+ cells, respectively, show comparable inhibition of tumor volumes at week 1. Remarkably, tumor volumes continue to shrink under these protocols even after treatment has ceased, with maximum reduction in volume predicted for strategy D. This behavior can be understood in terms of the long half-life of extracellular TMZ (48 h, estimated from [46]) and the time spent by DNA damaged cells in an arrested state and consequent delay in their apoptosis. However, by the end of the treatment cycle, xenografts treated with strategy B have recovered to attain their maximum possible volumes. In the long-term, only strategies C and D are predicted to have a significant impact on tumor growth with strategy D, outperforming all others by week 4. Thus, in the absence of any information regarding repair enzyme expression by tumor cells, we propose strategy D as the optimal protocol when treating heteroclonal tumors.

### In a virtual human clinical trial, strategy D predicts a 30% improvement in patient survival

Finally, we predict the potential survival benefit of treating human GBM patients with the identified optimal combination of TMZ and APNG/MGMT inhibitors (strategy D) by conducting a clinical trial *in silico*. Briefly, 100 virtual GBM patients were created, with each cancer comprised of a heterogeneous population of cells. We remark that in this case the range of proliferation rate was adjusted so that the mean doubling time was 49.6 days [47]. Each patient was treated with standard TMZ, optimal TMZ, and combination strategy D. We remark that were this an actual clinical trial, the patients would be randomized into one of the three treatment protocols, and their progression monitored. However, the *in silico* experimentation proposed here has the advantage that a virtual patient may be treated with as many strategies as desired. Thus, response to treatment may be compared without having to account for the confounding factor of inter-patient variability. Treatment was initiated once tumors reached a size of 1 cm in diameter (corresponding to 5 10^8^ cells), the average size of GBMs at diagnosis in a clinical setting [34], and continued for a maximum of 7 28-day cycles. This follows typical clinical protocol wherein TMZ is administered once (concomitant with focal radiotherapy, not considered here) followed by 6 cycles of TMZ alone [48]. Simulations were carried out until patient ‘death’, which was assumed to occur once tumors reached a critical size. This size was determined by the observation that, left untreated, GBM patients die within 3 to 4 months [49]. Details of modifications to the model necessary to simulate GBMs growing *in situ* are presented in the Supplemental Information.

A Kaplan-Meier survival analysis was conducted on our simulations, the results of which are summarized in figure 6. Under standard and optimal TMZ monotherapy, patient survival is predicted to decrease similarly (figure 6A) with comparable mean survival times of about 8 months. In particular, optimal TMZ improves mean survival by only one week as compared to standard TMZ. In contrast, under treatment with strategy D, all patients are expected to survive – barring adverse events not considered in our model – for up to 3 months after cessation of treatment. The mean survival under strategy D is over 10 months, a ~ 30% improvement over treatment with TMZ alone. A steep decline in survival is predicted thereafter (figure 6A, inset), due to the rapid growth of tumor cells left unaffected by therapy administration.

**Figure 6.**
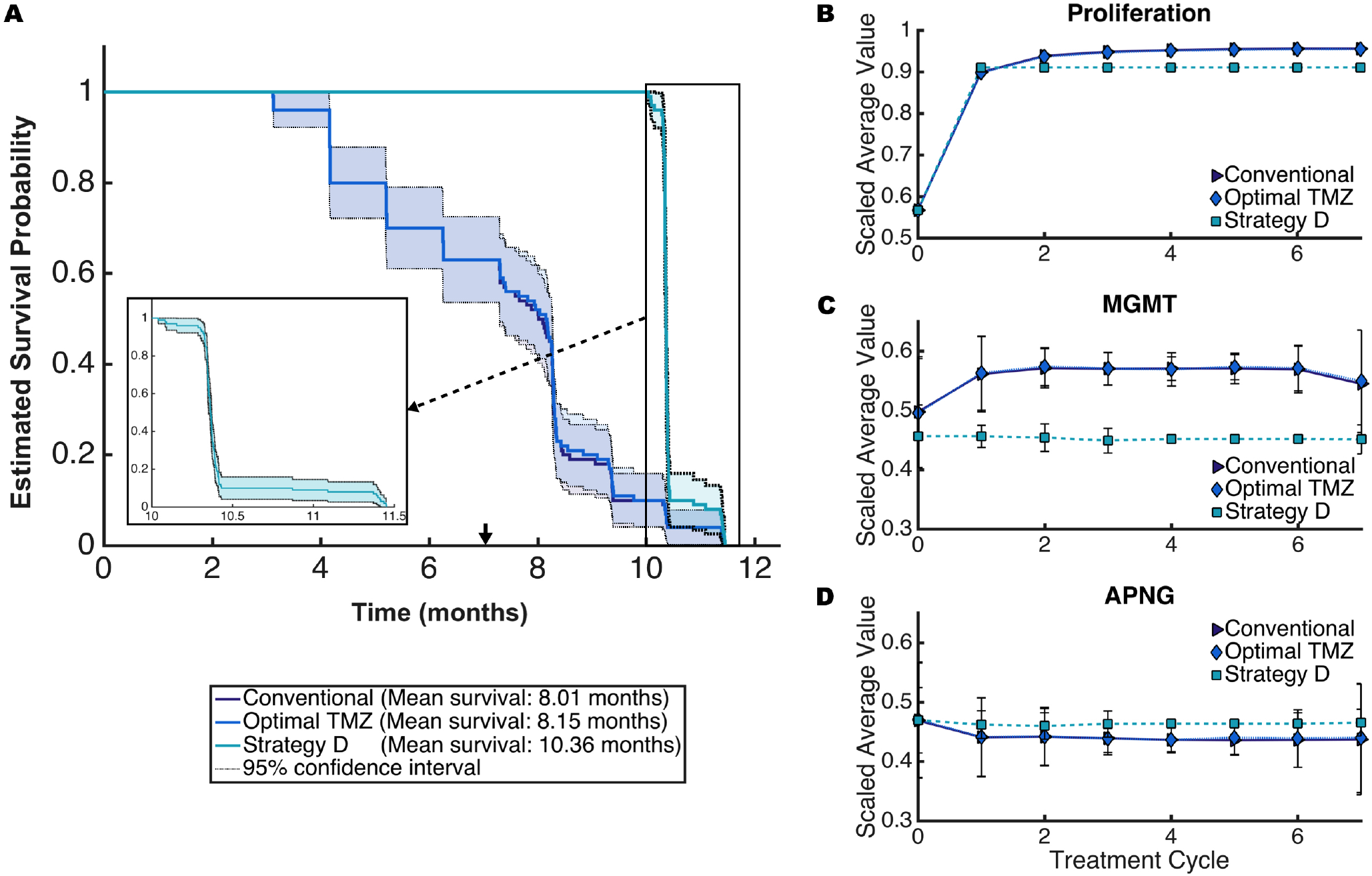
Results of an *in silico* human trial. A, Kaplan-Meier survival probabilities when patients were treated with standard TMZ, optimal TMZ, and combination strategy D for 7 28-days treatment cycles. B-D, proliferation rates, and APNG and MGMT expression averaged over each tumor over the course of 7 treatment cycles. Data points represent the mean across the 100 patients, with 95% confidence intervals.

With the development of targeted therapeutics such as those considered here, an important question is: "How do tumors evolve under the selection pressures created by the treatment?". To answer this, we recorded the composition of each patient’s tumor periodically, under each treatment strategy. Given the diversity in cell lines across all patients, the data was first quantile-normalized to make the distributions identical in statistical properties [50], and the resulting parameter distributions scaled by their respective maxima. The results are shown in figure 6B-D, which plot time-courses of the mean values and 95% confidence intervals of the scaled average tumor compositions.

Standard and optimal TMZ monotherapy exert similar selection pressures, resulting in tumors that have an overall bias towards cell lines with faster doubling times and high MGMT expression. Cellular investment in MGMT expression is compensated for by expressing lesser APNG, as observed in the case of heteroclonal xenograft response to TMZ (figure 3A-C). In particular, TMZ administration induces death in slow proliferating cells with low repair enzyme expression, leaving an abundance of space and resources for those cells over-expressing MGMT. Subsequent cycles of TMZ become less effective since they are targeting a population of cells capable of efficient DNA damage repair, underscoring the role played by MGMT in mediating TMZ-resistance. The emergent tumor phenotype is markedly different under strategy D, which favors cell lines that invest less in proliferation, less in MGMT expression, but more in APNG. Further, the emergent phenotype has very narrow 95% confidence intervals, indicating that tumors evolve to a more homogeneous phenotype under strategy D. This could explain the steep decline in patient survival after therapy cessation (figure 6A, inset).

## Discussion

GBM lethality is driven, in part, by resistance to TMZ, the most commonly used chemotherapeutic drug for treating this disease [4]. TMZ-induced DNA damage is efficiently repaired by the APNG-mediated BER repair pathway or by the MGMT pathway. Therefore, it has been hypothesized that combining TMZ with novel small molecule inhibitors of APNG and/or MGMT may improve patient survival. Indeed, several such drugs are in various stages of clinical trials [7, 15]. The mechanistic models studied here provide counter-intuitive results which could be crucial in the success or failure of TMZ-adjuvant clinical trials. First, in an emergent result, optimal scheduling across tumor phenotypes includes continuous inhibition of APNG-mediated DNA repair, even though MGMT repairs damage which is more lethal to the cancer cell. Second, only in MGMT over-expressing cancers is continuous MGMT-inhibition optimal, whereas even in tumors with low APNG expression, APNG inhibition is vital to cell killing potential.

The mechanistic modeling approach employed here is central to the predictive power of the models. Crucially, this model includes TMZ-induced DNA methylation and its subsequent repair. Our model was extensively parameterized and validated with available experimental data, resulting in a computational framework ideally suited to testing the anti-cancer potential of various drug combinations. GBM cells are known to vary highly at a molecular level, even within the same tumor [51]. Therefore, we needed to generate a phenotypically diverse population of GBM cell lines that is representative of different potential responses to treatment. For this, sensitivity analyses (summarized in figure 2) were performed on model parameters revealing rates of cellular proliferation and APNG and MGMT expression as critical drivers of TMZ-resistance. Pre-existing variation in the simulated cell populations was subject to constraints reflecting limited cellular capacity to both replicate rapidly and repair DNA damage with high fidelity. Over the course of these simulated experiments, selection on replicative potential and DNA-repair capacity of cell lines affected the proportions of different clones. Simulations of the response to treatment of polyclonal xenografts further elucidated determinants of resistance to TMZ (figure 3). In these single-dose studies, repair enzyme expression is found to affect cell survival more strongly than doubling time, consistent with the literature [10, 11].

Next, cell growth inhibition assays were simulated and a genetic algorithm employed to arrive at optimal dosing and scheduling protocols for each virtual cell line when TMZ was administered together with APNG and MGMT inhibitors. The protocols thus obtained revealed a natural grouping based on cell phenotype as determined by expression levels of APNG and MGMT. In all cases, alternating the dosing of TMZ, with high doses administered on days 1, 3 and 5, and low doses on days 2 and 4, was predicted to be optimal. Indeed, there is experimental evidence that cells over-expressing MGMT are more susceptible to such a schedule, which may have the additional advantage of reducing dose-limiting hematologic toxicity [45]. Against our expectations, APNG inhibition was necessary for maximizing cell kill even in experiments lacking high-APNG expressing subclones.

Finally, in a virtual pre-clinical trial, strategies optimal for cell lines over-expressing MGMT were predicted to be most successful in treating heteroclonal tumor xenografts when the overall phenotype of the cancer is unknown. We identified the strategy optimal for cells over-expressing both MGMT and APNG (strategy D) as the protocol with maximum tumor inhibition potential. A virtual clinical trial was then conducted wherein patients were enrolled in 3 treatment arms: standard TMZ monotherapy; alternating TMZ monotherapy; and strategy D combination therapy. Kaplan-Meier estimates showed that patients treated with strategy D continue to survive up to 3 months after treatment has ceased, with an overall improvement of 30% in mean survival time compared to those treated with TMZ alone. Both TMZ monotherapies had similar predicted survival curves, and therefore, the benefit of alternating TMZ may simply be mitigating side-effects [45]. We note that our model ignores adverse events in patients which may impact predicted survival times.

### Future experiments in cell-based and animal systems

The mathematical model presented here makes testable predictions on the effect of drug timing in cell lines and in animal models. In these experimental systems, biotechnology techniques such as RNAi [52] can be used to inhibit DNA repair enzymes even when drugs are not available; and, pre-clinical drugs can be delivered directly to cells even when highly toxic or lacking in bioavailability [53]. Using such approaches, predicted optimal drug combinations and schedules may be tested *in vitro* by replicating some of the virtual cell growth inhibition assays simulated here. Following the protocol in [41], cell cultures of human GBM cell lines could be established with a range of APNG or MGMT expression. These cell lines could be treated using different schedules of APNG inhibition, MGMT inhibition and TMZ corresponding to one of the four optimal strategies described above, with cell number tracked over the course of weeks. These experiments could confirm the *in silico* predictions made here, which would support the biological models of DNA repair and chemotherapy that these models encode. Further, experimental data thus generated would enable fine tuning of mechanistic model parameters and relaxing of modeling assumptions. Utilizing high-dimensional technologies such as luminex-bead expression assays [54], additional drug targets and potential adjuvant drugs could be identified. While such high-dimensional (“omics”) experiments are outside the mechanistic scope of the models presented here, an “omics” approach to studying relative drug timings, such as DIGRE [55, 56] becomes far more practical when a finite combination of administrations and timings can be proposed.

Independent of our model predictions, animal models should be used to assess for synergistic hematologic toxicity [45]. The mechanistic models reported here do not address the ratio of GBM-killing to hematologic toxicity of TMZ plus adjuvant therapy. Experimental results would be of tremendous utility in developing mechanistic models that might identify or predict such adverse synergistic effects.

### Potential for clinical translation

This model also makes testable predictions that could inform either clinical trials or expanded access (formerly known as “compassionate use”) of TMZ with adjuvants. There are several clinical trials and studies incorporating inhibition of MGMT, but fewer clinical trials inhibiting APNG [57, 58]. While MGMT plays a bigger role than APNG in chemotherapy resistance, we found that, in all cases, optimal treatment always includes APNG inhibitors. At the time of writing, we are unable to find trials that combine both inhibitors with TMZ.

When such trials are conducted, the timing and administration of novel drugs could mark the difference between a successful or failed trial or expanded access intervention. Randomized protocols for TMZ and bortezomib are actively recruiting patients (NIH clinical trial number NCT03643549). This trial incorporates an experimentally-motivated schedule, but experimental approaches cannot examine all possible schedules as is done here. Therefore, these trials are not utilizing the optimal schedule identified here. Expanded access would be justified by the lack of long-term effective treatments for this lethal disease [59], and could employ our optimal schedule. Even so, one limitation of our modeling approach should be emphasized: our model incorporates synergistic effectiveness between TMZ and MGMT/APNG inhibition, but does not include synergistic toxicity. Further, our model assumes that pharmacological inhibition of MGMT or APNG is total, while compounds like the MGMT-inhibitor bortezomib only achieve partial inhibition in a clinical setting [60].

### Conclusions

In both experimental and clinical contexts, the development and testing of novel treatment strategies in cancer is associated with a steep financial cost and significant human resource investment. Educated predictions about drug doses and schedules most likely to succeed are therefore critical for optimizing experimental design and resource investments on the part of the clinical investigator. *In silico* experimentation, such as that proposed here, affords the unique opportunity of simulating various treatment protocols over a wide range of parameters values that is not possible in a clinical setting. In our model of TMZ treated GBM, these simulations give an emergent result which supports both a novel combination therapy targeting APNG, and an intermittent schedule of TMZ.

## Data accessibility

Data sets and code is publicly available at (link will be provided). Code to perform identifiability analysis with GENSSI 2.0 was taken from https://github.com/genssi-developer/GenSSI. Code to use eFAST was taken from http://malthus.micro.med.umich.edu/lab/usadata/.

## Acknowledgements

We are grateful to Prof. Giray Okten, Mr. Nicholas Yaeger and Ms. Mayassa Dargham for many helpful discussions.

## Author Contributions

I.C.S. and H.V.J. conceived the experiments, I.C.S. conducted the experiments, I.C.S., S.K.H. and H.V.J. analyzed the results. All authors reviewed the manuscript.

