## Supplemental Material for "Mitigating Temozolomide Resistance in Glioblastoma via DNA Damage-Repair Inhibition"

### S1 Structural Identifiability

The specific dynamics of the model determine structural identifiability, which is considered to be an analytical property of the model independent of clinical data. In general, a lack of structural identifiability can only be addressed by reformulating the model and reducing the number of parameters. Such a step is crucial since most numerical approaches will be unable to estimate non-identifiable parameters accurately [1].

The general idea behind structural identifiability is addressing whether the model parameters can be uniquely estimated given a perfect collection of data on the model variables. Commonly used methods to determine structural identifiability include the Taylor series approach and the generating series method. We refer to [1] for a brief review of these and other methods. Here, we use the Matlab toolbox GenSSI 2.0, a multi-experiment structural identifiability analysis toolbox. It is the first toolbox for structural identifiability analysis to implement Systems Biology Markup Language import, state parameter transformations, and multi-experiment structural identifiability analysis [2, 3]. GenSSI 2.0 uses a generating series approach to determine identifiability, and the results are presented in identifiability tableaus.

The generating series approach establishes a system of non-linear algebraic equations where the observables are expanded in series with respect to time and inputs by calculating subsequent derivatives. The coefficients of such series are the output functions and their successive Lie derivatives [1, 2, 3]. These Lie derivatives are evaluated at $t = t_0$, put into vector equation form. Using the inverse function theorem, this vector equation can be uniquely solved (locally) if and only if the Jacobian matrix has full rank. This is known as the rank-test introduced by Pohjanpalo in 1978 [4].

One disadvantage of this method is that the minimum number of Lie derivatives needed is unknown, and the process must be done with an initial guess. This can result in several iterations before obtaining results, especially using GenSSI 2.0. It is worth noting that, although conceptually simple, this approach may generate algebraic equations that may be challenging or impossible to solve, even with the aid of a symbolic manipulation software [1].

Identifiability tableaus are a powerful tool that can help overcome some of the limitations by providing information about the relationship between parameters. They represent the non-zero elements of the Jacobian of the series of coefficients with respect to the parameters, as explained in [5]. Each tableau has as many columns as parameters are studied and as many rows as non-zero series coefficients. A black square in the coordinate $(i, j)$ indicates that the non-zero generating series coefficient $i$ depends on the $j^{th}$ parameter [1]. Parameters may be unidentifiable when the Jacobian rank is deficient, presenting empty columns. When the rank of the Jacobian is full (i.e., coincides with the number of parameters) all parameters are, at least, locally identifiable. The analysis has three possible outcomes: globally identifiable, that is the parameter could be found no matter the range of the parameters; locally identifiable, that is the uniqueness of the parameters can only be determined within a neighborhood of parameter values, not all the possible parameter values; and unidentifiable, that is the parameter does not have a unique solution [2, 3].

In the case when the rank of the Jacobian is equal to the number of parameters, identifiability tableaus are particularly useful. Usually, the number of non-zero coefficients is larger than the number of non-zero parameters. This means that we can generate a minimal square tableau by choosing rows that guarantee a full rank in the Jacobian. In such minimal tableaus if a parameter is the unique non-zero element in a particular row, then the parameter is structurally identifiable. In that case, we calculate the parameter as a function of the power series coefficients and eliminate it to create a reduced tableau. Once all the possible reductions are made, we solve the remaining equations. The tableu also helps determine what equations are necessary to solve for each parameter, since it is not often the case where all the power series depend on all parameters.

Our proposed model is divided into the sub-cellular and the population scale. Since clinical data needed to estimate the parameters in each of these scales comes from various sources, and in each case the model needs to be reduced to be consistent with the experiments set up, we do not consider the full model for structural identifiability of all parameters, but rather we performed the study in each reduced model. We first performed identifiability analysis in equations (**??**), the system of equations for DNA damage. We remark that the PK/PD structure in the intracellular-extracellular transport of TMZ renders the model unidentifiable. Unidentifiability arises from the volume distributions $V_o$ and $V_i$. This phenomenon is a well-known result for these types of models, where the volume distributions need to be prescribed. Once these two parameters were fixed for the analysis, four Lie derivatives guaranteed full Jacobian rank (six), thus deeming all parameters at least structurally locally identifiable. Upon further analysis, we determined that the parameters $\gamma_{oi}, \gamma_{io}$, and $K_d$ are structurally globally identifiable, while the rest of the parameters ($k_f^N, k_f^O, k_r^N, k_r^O, k_p^N$, and $k_p^O$) are locally identifiable. However, given the biological nature of our model, we do not need to change the structure of the model since we will be able to provide a range for these parameters' values *a priori*. Identifiability tableaus for this analysis are presented in figure S1A-E. Similarly, an analysis of equations (**??**), the system of equations for DNA repair, was performed. In this case, we achieved the rank of the full Jacobian, ten, with only two Lie derivatives. All the parameters ($k_f^A, k_f^M, k_r^A, k_r^M, k_p^A, k_p^M, S_{APNG}, S_{MGMT}, \lambda_{APNG}$, and $\lambda_{MGMT}$) are structurally globally identifiable. Identifiability tableaus for this analysis are presented in figure S1F-H.

Finally, we performed the analysis in the population model equations (**??**), where a discretization of the age-structured was needed to use the generating series approach. After eight Lie derivatives, the rank of the Jacobian was full and local identifiability was guaranteed for all parameters. In this case, $\alpha, K, \mu$, and $\rho$ were determined to be globally identifiable. However, $a_0, b_1, b_2, c_1, c_2$, and $c_3$ are simply locally identifiable.

An important conclusion from the analysis is that, if $a_0 = 0$, losing the age-structured in arrested cells, structural identifiability of the model is no longer achievable. Therefore, we claim that the age-structure is essential not only to capture biological detail but also for mathematical well-posedness.

### S2 Parameter Estimation

The following parameters are found by fitting to available clinical data: proliferation rate ($\alpha$) and carrying capacity ($K$) of tumor cells; enzymatic production and degradation ($S_{APNG}, S_{MGMT}, \lambda_{APNG}, \lambda_{MGMT}$); rates constants related to DNA methylation ($k_f^N, k_r^N, k_p^N, k_f^O, k_r^O, k_p^O$) and DNA repair ($k_f^A, k_r^A, k_p^A, k_f^M, k_r^M, k_p^M$); parameters related drug transport in/out of cells ($V_{out}, V_{in}, \gamma_{oi}, \gamma_{io}, K_d$); and parameters governing cell arrest ($\mu$), cell repair ($\rho$), and cell death ($b_1, b_2, c_1, c_2, c_3$). Clinical data is obtained from a variety of sources in the literature, hence for each series of experiments, the model is simplified to fit the conditions the experiments were carried out in. We now present a description of this process.

*Tumor Growth ($\alpha, K$)* – In a series of experiments described in [6, 7, 8, 9, 10], GBM cell lines A172, U251, C6, and T98G were cultured and recorded periodically. Cell lines A172 were plated for 6 days in modified Eagle's medium containing 406 mg/l L-alanyl-L-glutamine, 10% fetal calf serum, 100 $\mu$g/ml penicillin, and 100 $\mu$g/ml streptomycin [6, 7]. Cell lines U251 were plated and serum-starved for one day, with an initial count of $0.75 \times 10^4$ cells [8]. Cell lines C6 were seeded into ten dishes, 35 mm in diameter, with an initial count of $5 \times 10^4$ cells [9]. Cell lines T98G were seeded in triplicate in tissue culture dishes with an initial count of $0.2 \times 10^6$ cells [10]. These data were used to fit $\alpha$ and $K$ in the reduced equation of proliferating cells

$$\frac{dP}{dt} = \alpha P(t)\left(1 - \frac{P(t)}{K}\right). \tag{S1}$$

Best fits are shown in figure S2A, with the parameter value estimates recorded in table S3.

*Repair Enzymes half life* – The half-life of MGMT in the presence of methylating agents is approximately 160 min [11], so that $\lambda_{MGMT} = 0.0043$ min$^{-1}$. The half-life of APNG is approximately 120 min [12], thus $\lambda_{APNG} = 0.00577$ min$^{-1}$.

*DNA Damage Rates ($k_f^N, k_r^N, k_p^N, k_f^O, k_r^O, k_p^O$)* – In a series of experiments described in [13] Big Blue Rat2 cells were treated with TMZ to investigate the rates of DNA methylation. Briefly, Big Blue Rat2 cells were grown in 175 cm$^2$ flasks containing 25 ml of medium. Various amounts of TMZ were added directly to the culture medium, and after three hours the cells were washed. DNA was isolated from the cellular pellets, and the levels of N7-meG and O6-meG adducts were collected. Figures S2B and C show these data as a function of drug added. In the same experiments, time course data of adduct formation was also recorded. DNA was incubated with 500$\mu$M TMZ for various lengths of time at 37$^o$C. Figure S2D shows the corresponding time courses. In all these experiments DNA was isolated from the cells; thus a population model is not valid to estimate these parameters. We also neglect intracellular transport of TMZ in/out of cells since DNA was not collected until TMZ concentration was homogeneous everywhere. Lastly, because the cells were washed after only a few hours of TMZ exposure, we assume that DNA damage repair would not have been initiated. These data were used to estimate $k_f^N, k_r^N, k_p^N, k_f^O, k_r^O, k_p^O$ in the reduced model

$$\begin{aligned}
\frac{dT_{in}}{dt} &= -k_f^N N_7 T_{in} - k_f^O O_6 T_{in} + k_r^N D_7 + k_r^O D_6,\\
\frac{dN_7}{dt} &= -k_f^N N_7 T_{in} + k_r^N D_7,\\
\frac{dD_7}{dt} &= k_f^N N_7 T_{in} - k_r^N D_7 - k_p^N D_7,\\
\frac{dA_7}{dt} &= k_p^N D_7,\\
\frac{dO_6}{dt} &= -k_f^O O_6 T_{in} + k_r^O D_6,\\
\frac{dD_6}{dt} &= k_f^O O_6 T_{in} - k_r^O D_6 - k_p^O D_6,\\
\frac{dA_6}{dt} &= k_p^N D_6.
\end{aligned} \tag{S2}$$

Best fits are shown in S2B-D, with the parameter value estimates recorded in table S4.

*TMZ Transport in/out of Cells ($\gamma_{oi}, \gamma_{io}, K_d$)* – In a series of experiments described in [14], human GBM cells were exposed to TMZ to study the transport of the drug in an out of the cell. A total of 106 U87 cells were seeded onto 60 mm Petri dishes and cultured overnight. Next, 50 $\mu$mol/l of TMZ was added, and intracellular and extracellular concentration of TMZ were sampled for two hours. These data were used to estimate $\gamma_{oi}, \gamma_{io}$, and $K_d$, as well as the initial values of DNA that can be methylated in the N7 and O6 positions in the reduced equations:

$$\begin{aligned}
\frac{dT_{out}}{dt} &= -\gamma_{oi} T_{out} + \gamma_{io} \frac{V_{in}}{V_{out}} T_{in} - K_d T_{out},\\
\frac{dT_{in}}{dt} &= \gamma_{oi} \frac{V_{out}}{V_{in}} T_{out} - \gamma_{io} T_{in}.
\end{aligned} \tag{S3}$$

Best fits for extracellular and intracellular TMZ concentrations are shown in figure S2E, and F (blue and red lines respectively), and parameter values in table S4. Volumes of distribution were taken from [14].

*DNA Repair Rates ($k_f^A, k_r^A, k_p^A, k_f^M, k_r^M, k_p^M$)* – In a series of experiments described in [15] clinical data of the MGMT repair process was obtained. DNA containing O6-meG oligonucleotides was incubated with different concentrations of MGMT. Reactions were initiated by rapid mixing of MGMT (12.5 $\mu$l) with DNA substrate (10.9 $\mu$l). The mixtures were incubated at 37$^o$C for 30 min. Demethylation rate of O6-meG DNA for different concentrations of MGMT was recorded and are shown in figure S2D (blue circles). These data were used to estimate the parameters in the reduced equations

$$\begin{aligned}
\frac{dO_6^*}{da} &= k_p^M C_6^*,\\
\frac{dA_6^*}{da} &= -k_f^M A_6^* MGMT + k_r^M C_6^*,\\
\frac{dC_6^*}{da} &= k_f^M A_6^* MGMT - k_r^M C_6^* - k_p^M C_6^*,\\
\frac{dMGMT}{da} &= S_{MGMT} - \lambda_{MGMT} MGMT - k_f^M A_6^* MGMT + k_r^M C_6^*.
\end{aligned} \tag{S4}$$

We consider $C_6^*$ to be at quasi-steady state assuming Michaelis-Menten kinetics, and since the total amount of DNA should be conserved while the repair process is happening we can further simplify our model by taking $O_6^* + A_6^* + C_6^* = DNA(0)$ a constant. Under these assumptions a functional solution for $C_6^*(a)$ and $A_6^*(a)$ reads

$$\begin{aligned}
K^M &= \frac{k_r^M + k_p^M}{k_f^M},\\
C_6^*(a) &= \frac{(DNA(0) - O_6^* - A_6^*) MGMT}{K^M + MGMT},\\
A_6^*(a) &= \frac{(DNA(0) - O_6^* - C_6^*) K^M}{K^M + MGMT},
\end{aligned} \tag{S5}$$

and the velocity of O6-meG demethylation is given by

$$v_{O_6}(a) = \frac{k_p^M MGMT}{K^M + MGMT}. \tag{S6}$$

Best fist are shown in figure S2D, and estimations for $k_p^M$ and $K_M$ are redorded in table S4. Likewise, the velocity of DNA repair by APNG can be derived from the reduced model

$$\begin{aligned}
\frac{dN_7^*}{da} &= k_p^A C_7^*, \\
\frac{dA_7^*}{da} &= -k_f^A A_7^* APNG + k_r^A C_7^*, \\
\frac{dC_7^*}{da} &= k_f^A A_7^* APNG - k_r^A C_7^* - k_p^A C_7^*, \\
\frac{dAPNG}{da} &= S_{APNG} - \lambda_{APNG} APNG - k_f^A A_7^* APNG + k_r^A C_7^*,
\end{aligned} \tag{S7}$$

by taking $C_7^*$ to be at quasi-steady state, which under the previous assumptions resolves

$$\begin{aligned}
K^A &= \frac{k_r^A + k_p^A}{k_f^A}, \\
C_7^*(a) &= \frac{(DNA(0) - N_7^* - A_7^*)APNG}{K^A + APNG}, \\
A_7^*(a) &= \frac{(DNA(0) - N_7^* - C_7^*)K^A}{K^A + A},
\end{aligned} \tag{S8}$$

which yields

$$v_{N_7}(a) = \frac{k_p^A APNG}{K^A + APNG}. \tag{S9}$$

Estimates of $k_p^A$ and $K^A$ are available in [16].

*Damage in vitro ($\mu, \rho, b_1, b_2, c_1, c_2, c_3$)* – In a series of experiments described in [17], the human cell line A172 was used to investigate whether APNG and MGMT confers TMZ resistance. A172 are human GBM cells that do not express MGMT nor APNG; thus, they are an optimal choice to understand how APNG and MGMT expression affects cell viability. The cells were transfected with constructs containing APNG, MGMT, or both, and incubated with various concentrations of TMZ ($0 - 250$ $\mu$M) to conduct survival assays. A total of $1 \times 10^5$ cells were plated into six-well dishes in 2 ml. After incubation with various concentrations of TMZ, cells were collected and analyzed. This data was used to estimate the parameters related to cell arrest, recovery, and cell death $\mu, \rho, b_1, b_2, c_1, c_2$, and $c_3$, and the full model was used. Best fits are shown in figure S3B, and parameter estimations are recorded in table S4.

### S3 Parameter Validation

Following the experiments described in [17], cell viability assays were performed with different GBM cell lines that are known to express various levels of APNG and MGMT. The cell lines studied were: T98G expression both APNG and MGMT, A172 cells expressing neither, C6 rat glioma cells which express predominantly MGMT and moderate to low APNG, and U251 human GBM cells which express APNG but not MGMT. Using appropriate values of $\alpha$ and $K$ for each cell line we conducted model simulations treating cell lines with $0 - 250$ $\mu$M of TMZ, and recorded cell viability. Clinical data along with such model simulations are shown in figure S3. The model simulations show good agreement with this data, validating our model formulation.

### S4 Energy Inequality

We derive an energy constraint inequality to which all cells must adhere motivated by the approach of Nagy et al. [18]. They introduced an energy model with the view to explaining the angiogenic switch in cancer with a multi-scale model that includes an energy management system in an evolving angiogenic tumor.

We consider tumor cells to divide their efforts into maintaining normal physiological function, proliferating, and production of repair enzymes. The total maintenance rate is taken to be $\eta_\beta = 8$ min$^{-1}$, and the proliferation effort to be $\eta_P = 3.5$ min$^{-1}$ (as described in section 4.3 in [18]). Additionally, we consider the efforts to produce APNG ($\eta_A$) and MGMT ($\eta_M$). Tumor cells are assumed to grow at their maximum rate burning adenosine triphosphate (ATP) at a rate of approximately 22 fmol min$^{-1}$ while maintaining their ATP level at 1.5 fmol:

$$1.5(\eta_\beta + \eta_P + \eta_M + \eta_A) \approx 22. \tag{S10}$$

Thus, assuming repairing both types of DNA damage is equally expensive, we can calculate that $\eta_A = \eta_M = 1.58$ min$^{-1}$.

Now, we can think of cellular energy in terms of production of cellular material such as APNG, MGMT, or any cellular material needed for cell division. In such terms, we define energy as the effort required to create 1 $\mu$M of cellular material in 1 min, where effort related to the rate of ATP consumption. We assume that total cellular energy available is kept constant in all cells and that it can be decomposed as

$$E_C = \eta_P S_P + \eta_A S_{APNG} + \eta_M S_{MGMT} + \eta_\beta S_\beta. \tag{S11}$$

Where $S_P$ is the production of cellular material, $S_{APNG}$ is the production of APNG, $S_{MGMT}$ is the production of MGMT, and $S_\beta$ is the production of any cellular material needed to maintain normal physiological function. Further we consider

$$S_P = \alpha x, \tag{S12}$$

Where $\alpha$ is the proliferation rate of tumor cells, and $x$ is a characteristic concentration of cellular material, that for simplicity we take to be $x = 1$ $\mu$M. Moreover, $S_\beta$ is considered to be constant for all cell lines. Under these assumptions, we propose an upper bound for $E_C$, $M_{E_C}$ given by the cellular energy of T98G cells, an established GBM cell line known for its virulence. Thus all virtual cell lines must satisfy the constraint

$$\eta_P S_P + \eta_A S_{APNG} + \eta_M S_{MGMT} \leq 2373.46. \tag{S13}$$

We create multiple virtual cohorts of tumor cell lines where $\alpha, S_{APNG}$, and $S_{MGMT}$ are randomly selected in such a way that equation (S13) is satisfied. The three studied cohorts are as follows:

- **Monoclonal *in vitro* cohort**: 500 monoclonal tumors grown *in vitro*. Each tumor is comprised of a single cell type. This cohort is equally divided into four groups of cell lines depending on their expression of APNG and MGMT. The first group is characterized by low expressions of MGMT and APNG; the second by low expression of APNG and high of MGMT; the third by low expression of MGMT and high of APNG; and the last group by high expression of MGMT and APNG. This cohort resembles *in vitro* experiments done with established cell lines such as A172, U251, C6, and T98G.

- **Heteroclonal xenograft cohort**: 500 heteroclonal tumors grown in mice. Each tumor is comprised of 10 different cell lines. This cohort resembles xenograft experiments done with GBM samples from patients.

- **Heteroclonal human cohort**: 100 heteroclonal tumors grown in patients. Each tumor is comprised of 10 different cell lines. This cohort resembles GBM patients. Specifically, tumors are allowed to grow exponentially, without a carrying capacity $K$, because patients' death is assumed once the number of tumor cells reaches a certain threshold. Further, the range of $\alpha$ values is adjusted so that the mean doubling time is 49.6 days, matching clinical data of growth dynamics of untreated GBMs *in vivo* [19].

### S5 Genetic Algorithm Details

Genetic algorithms mimic the principles of genetic evolution to find the solution to an optimization problem, in our case, tumor response to treatment. An initial pool of treatment strategies is created randomly. The treatments are sorted based on their fitness which is measured in terms of cell survival post-treatment. The top 20% of strategies are paired to create offspring. As in evolution, the offspring strategies inherit different properties from each of their parents. The resulting offspring are also allowed a small chance of mutation, thus creating a new generation of treatment protocols that are sorted again based on fitness, and the process repeated until convergence.

Genetic algorithms have been shown to converge rapidly to optimal solutions of problems that have multiple constraints, are irregular, or even discontinuous, which are difficult to solve by other traditional optimization methods. They implement multidirectional search, explore the search space using stochastic processes rather than deterministic rules, and are very flexible in the choice of the function that is being optimized [20]. We implemented our genetic algorithm using Julia, setting the selection rate to be 20%, mutation rate to be 5%, an initial population of 20 treatment strategies, and a total number of iterations of 20, since consistent results were achieved past that number of iterations in initial studies. Each treatment strategy is assumed to be a 28-day cycle, where treatment is given in days 1 to 5 of week 1, following the current schedule of TMZ given to GBM patients. Each day of treatment three decisions are made: TMZ dose, administration of MGMT inhibitor, APNG inhibitor, both, or neither. Thus a treatment strategy will comprise a varying dose of TMZ coupled with a schedule of APNG and MGMT inhibitors.

**Algorithm:** Genetic Algorithm for selecting treatment strategies

1. Create a set of 20 treatment strategies chosen randomly.
   (a) TMZ dose: Create a vector of length 5 selecting values randomly. Check that the total dose does not surpass the maximum dose.
   (b) MGMT inhibitor: Create a vector of length 5 with 0's and 1's chosen randomly.
   (c) APNG inhibitor: Create a vector of length 5 with 0's and 1's chosen randomly.
2. Treat the cell line under each of the 20 strategies.
3. Rank strategies based on the number of tumor cells after treatment.
4. Select the top 20% of strategies.
5. Randomly paired the selected strategies and create 20 offspring strategies:
   (a) TMZ dose: Create a vector of length 5 with 1's and 2's chosen randomly, $\mathbf{w}$. Now, create a vector of length 5 $\mathbf{v}$, for $i = 1,\ldots,5$, if $\mathbf{w}_i = 1$, then $\mathbf{v}_i$ is the same TMZ value as the parent strategy 1 had on day $i$. If $\mathbf{w}_i = 2$, then $\mathbf{v}_i$ is the same TMZ value parent strategy 2 had on day $i$. Once all the entries in the vector are selected, check that maximum dosage is not surpassed. If the total TMZ dosage is too high, repeat the process.
   (b) MGMT inhibitors: Create a vector of length 5 with 1's and 2's chosen randomly, $\mathbf{w}$. Now, create a vector of length 5 $\mathbf{v}$, for $i = 1,\ldots,5$, if $\mathbf{w}_i = 1$, then $\mathbf{v}_i$ is the same decision about MGMT inhibitor as the parent strategy 1 had on day $i$. If $\mathbf{w}_i = 2$, then $\mathbf{v}_i$ is the same TMZ value parent strategy 2 had on day $i$.
   (c) APNG inhibitors: Create a vector of length 5 with 1's and 2's chosen randomly, $\mathbf{w}$. Now, create a vector of length 5 $\mathbf{v}$, for $i = 1,\ldots,5$, if $\mathbf{w}_i = 1$, then $\mathbf{v}_i$ is the same decision about APNG inhibitor as the parent strategy 1 had on day $i$. If $\mathbf{w}_i = 2$, then $\mathbf{v}_i$ is the same TMZ value parent strategy 2 had on day $i$
6. Allow a small chance of mutation in the resulting strategies
7. Repeat from step 2.

| Name | Description | Units |
|---|---|---|
| $t$ | Time | min |
| $a$ | Time of Arrest | min |
| $P$ | Proliferating Cells | number of cells |
| $A$ | Arrested Cells | number of cells |
| $T_{out}$ | Extracellular Drug Concentration | $\mu$M |
| $T_{in}$ | Intracellular Drug Concentration | $\mu$M |
| $N_7$ | Healthy N7-guanines in proliferating cells | $\mu$M |
| $O_6$ | Healthy O6-guanines in proliferating cells | $\mu$M |
| $N_7^*$ | Healthy N7-guanines in arrested cells | $\mu$M |
| $O_6^*$ | Healthy O6-guanines in arrested cells | $\mu$M |
| $D_7$ | N7-guanine bound to TMZ | $\mu$M |
| $D_6$ | O6-guanine bound to TMZ | $\mu$M |
| $A_7$ | N7-methylguanine adducts in proliferating cells | $\mu$M |
| $A_6$ | O6-methylguanine adducts in proliferating cells | $\mu$M |
| $A_7^*$ | N7-methylguanine adducts in arrested cells | $\mu$M |
| $A_6^*$ | O6-methylguanine adducts in arrested cells | $\mu$M |
| $MGMT$ | MGMT repair enzyme | $\mu$M |
| $APNG$ | APNG repair enzyme | $\mu$M |
| $C_7^*$ | Repairing compound of N7-methylguanine and APNG | $\mu$M |
| $C_6^*$ | Repairing compound of O6-methylguanine and MGMT | $\mu$M |

**Table S1.** List of model variables with biological descriptions and units.

| Name | Description | Units |
|---|---|---|
| $k_f^N$ | Rate of N7-G and drug binding | $\text{min}^{-1}\ \mu\text{M}^{-1}$ |
| $k_r^N$ | Reverse rate of N7-G and drug binding | $\text{min}^{-1}$ |
| $k_p^N$ | Rate of N7-G methylation | $\text{min}^{-1}$ |
| $k_f^O$ | Rate of O6-G and drug binding | $\text{min}^{-1}\ \mu\text{M}^{-1}$ |
| $k_r^O$ | Reverse rate of O6-G and drug binding | $\text{min}^{-1}$ |
| $k_p^O$ | Rate of O6-G methylation | $\text{min}^{-1}$ |
| $k_f^A$ | Rate of N7-meG and APNG binding | $\text{min}^{-1}\ \mu\text{M}^{-1}$ |
| $k_r^A$ | Reverse rate of N7-meG and APNG binding | $\text{min}^{-1}$ |
| $k_p^A$ | Rate of N7-meG repair | $\text{min}^{-1}$ |
| $k_f^M$ | Rate of O6-meG and MGMT binding | $\text{min}^{-1}\ \mu\text{M}^{-1}$ |
| $k_r^M$ | Reverse rate of O6-meG and MGMT binding | $\text{min}^{-1}$ |
| $k_p^M$ | Rate of O6-meG repair | $\text{min}^{-1}$ |
| $V_{out}$ | Volume of extracellular drug distribution | ml |
| $V_{in}$ | Volume of intracellular drug distribution | ml |
| $K_d$ | Rate of degradation of drug | $\text{min}^{-1}$ |
| $\gamma_{oi}$ | Rate of TMZ entry into the cell | $\text{min}^{-1}$ |
| $\gamma_{io}$ | Rate of TMZ exit out of the cell | $\text{min}^{-1}$ |
| $S_{MGMT}$ | Production of MGMT | $\text{min}^{-1}\ \mu\text{M}$ |
| $S_{APNG}$ | Production of APNG | $\text{min}^{-1}\ \mu\text{M}$ |
| $\lambda_{MGMT}$ | Degradation of MGMT | $\text{min}^{-1}$ |
| $\lambda_{APNG}$ | Degradation of APNG | $\text{min}^{-1}$ |
| $\alpha$ | Proliferation rate of proliferating cells | $\text{min}^{-1}$ |
| $K$ | Carrying capacity of proliferating cells | number of cells |
| $\mu$ | Rate of cell arrest | $\text{min}^{-1}$ |
| $\rho$ | Rate of cell repair | $\text{min}^{-1}$ |
| $b_1$ | Rate of cell death caused by N7-meG damage | $\text{min}^{-1}$ |
| $b_2$ | Rate of cell death caused by O6-meG damage | $\text{min}^{-1}$ |
| $c_1$ | Rate of cell death caused by long arrest | $\text{min}^{-1}$ |
| $c_2$ | Effect of MGMT on cell death | $\mu\text{M}^{-1}$ |
| $c_3$ | Effect of N7-meG adducts on cell death | $\mu\text{M}^{-1}$ |
| $a_0$ | Characteristic waiting time of arrested cells | min |

**Table S2.** List of model parameters with biological descriptions and units.

| Cell Line | $\alpha$ (day$^{-1}$) | $K$ (number of cells) |
|---|---|---|
| A172 | 0.4535 | $5.8 \times 10^6$ |
| U251 | 0.8505 | $6.77 \times 10^6$ |
| T98G | 0.9913 | $11.6072 \times 10^6$ |
| C6 | 0.963 | $8.0483 \times 10^6$ |

**Table S3.** Tumor growth parameter estimations.

| Parameter | Value | Units | Reference |
|---|---|---|---|
| $\lambda_{MGMT}$ | 0.0043 | min$^{-1}$ | [11] |
| $\lambda_{APNG}$ | 0.0057 | min$^{-1}$ | [12] |
| $k_p^A$ | 0.35 | min$^{-1}$ | [16] |
| $K^A$ | 0.025 | $\mu$M | |
| $V_{out}$ | $2 \times 10^{-3}$ | l | [14] |
| $V_{in}$ | $7 \times 10^{-6}$ | l | |
| $k_f^N$ | 0.1270 | min$^{-1}$ $\mu$M$^{-1}$ | Fitted |
| $k_r^N$ | 0.0973 | min$^{-1}$ | |
| $k_p^N$ | 0.0130 | min$^{-1}$ | |
| $k_f^O$ | 0.5003 | min$^{-1}$ $\mu$M$^{-1}$ | |
| $k_r^O$ | 0.0457 | min$^{-1}$ | |
| $k_p^O$ | 0.0159 | min$^{-1}$ | |
| $k_p^M$ | 0.1717 | min$^{-1}$ | |
| $K^M$ | 0.0916 | $\mu$M | |
| $\gamma_{oi}$ | 0.0440 | min$^{-1}$ | |
| $\gamma_{io}$ | 29.086 | min$^{-1}$ | |
| $K_d$ | 0.0143 | min$^{-1}$ | |
| $\mu$ | 174.2872 | min$^{-1}$ | |
| $\rho$ | 2.9288 | min$^{-1}$ | |
| $b_1$ | 2.6411 | min$^{-1}$ | |
| $b_2$ | 7.2196 | min$^{-1}$ | |
| $c_1$ | 24.383 | min$^{-1}$ | |
| $c_2$ | 55.0068 | $\mu$M$^{-1}$ | |
| $c_3$ | 0.0112 | $\mu$M$^{-1}$ | |

**Table S4.** Other parameter estimations.

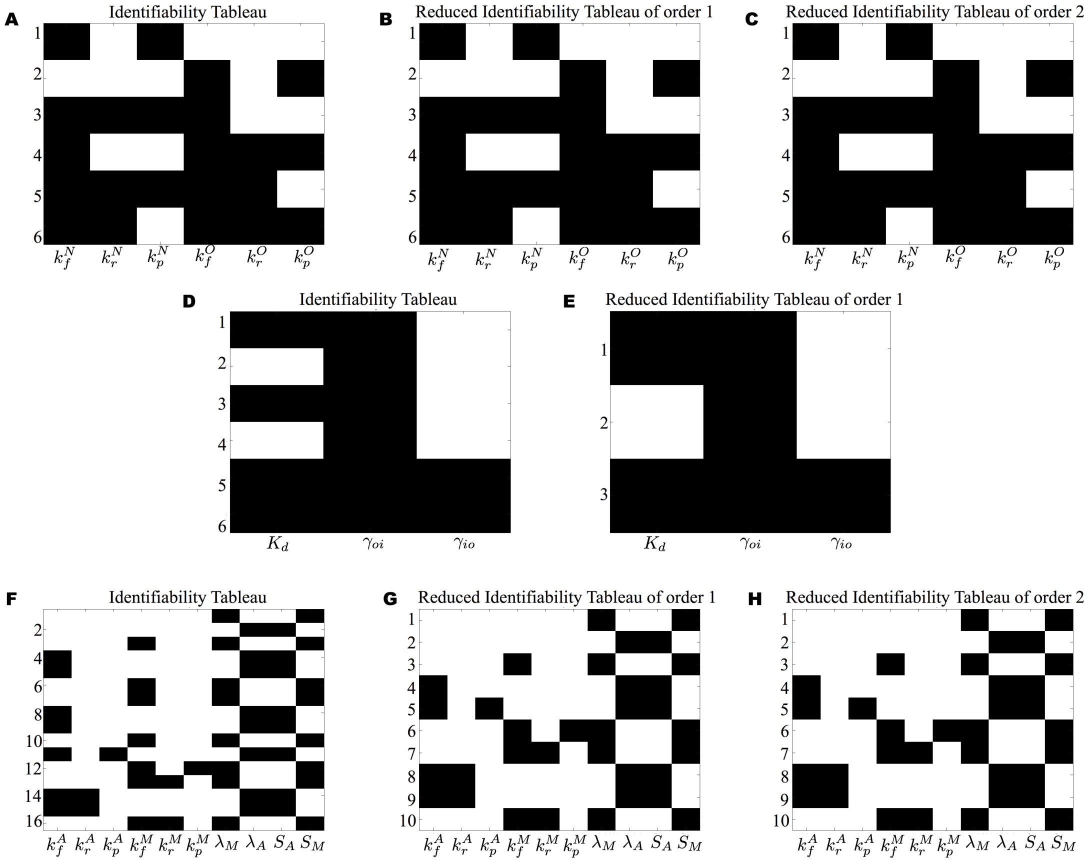


**Figure S1. Identifiability Tableaus.** Identifiability analysis was performed with GenSSI 2.0. The rows correspond to the generating series coefficients of the successive Lie derivatives of the model output. The columns correspond to the different parameters. A black square indicated that the corresponding non-zero generating series coefficient depends on the parameter corresponding to the column [1]. A-C Identifiability tableau, reduced identifiability tableau order 1, and reduced identifiability tableau order 2 for equations (**??**). The Jacobian matrix has rank 6. D-E Identifiability tableau, and reduced identifiability tableau order 1 for the intracellular PK/PD equations. The Jacobian matrix has rank 3. F-H Identifiability tableau, reduced identifiability tableau order 1, and reduced identifiability tableau order 2 of equations (**??**). The Jacobian matrix has rank 10.

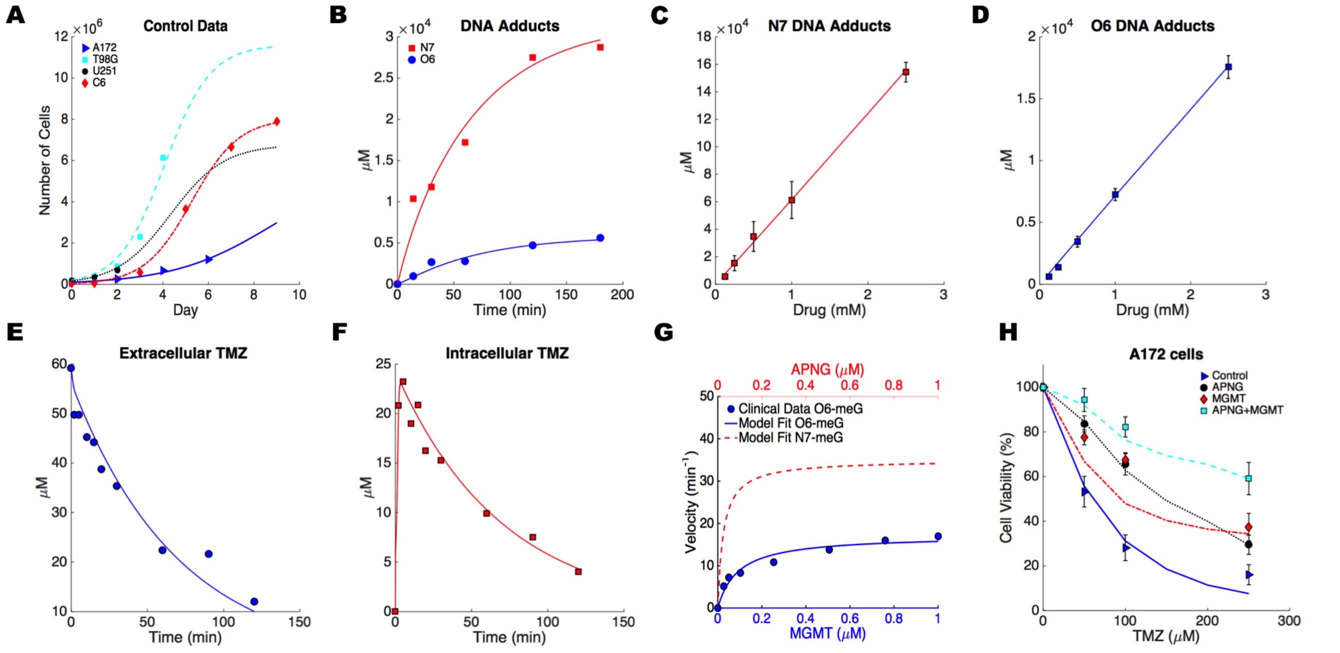


**Figure S2. Best fits to clinical data.** Clinical data used for parameter estimation is shown with the best fist of the model. In all cases, markers represent clinical data, and lines represent the model output. A Control data for cell lines A172, T98G, U251, and C6 was taken from multiple sources [6, 8, 9, 10]. Briefly between $0.75 \times 10^4$ and $2 \times 10^6$ cells were platted and harvested at different time points. B-D In a series of experiments described in [13] Big Blue Rat2 cells were treated with TMZ to investigate the rates of DNA methylation. B Time course of DNA adduct formation after 3h of treatment with 500 $\mu$M of TMZ. C Formation of N7-meG adducts as a function of TMZ. D Formation of O6-meG adducts as a function of TMZ. E-F In a series of experiments described in [14], human GBM cells were exposed to TMZ to study the transport of the drug in an out of the cell. E Time course of extracellular concentration of TMZ. F Time course of Intracellular concentration of TMZ. G In a series of experiments described in [15] clinical data of the MGMT repair process was obtained. DNA containing O6-meG oligonucleotides was incubated with different concentrations of MGMT. The velocity of O6-meG demethylation is shown (blue circles), along with model best fits. The velocity of N7-meG demethylation rate of N7-meG was obtained using parameter values reported in [16] and model simulations are shown in red. H In a series of experiments described in [17] A172 cells were transfected with constructs containing APNG, MGMT, or both, and incubated with various concentrations of TMZ. Cell viability of A172 was recorded in each case at the end of the experiment.

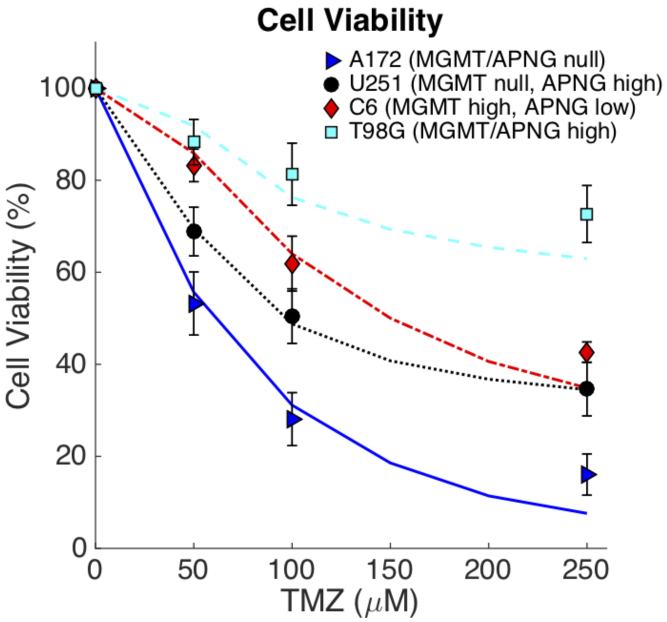


**Figure S3.** **Parameter Validation.** Clinical data of cell lines A172, U251, C6, and T98G treated with TMZ was used to validate the model parametrization. Briefly, cell lines were incubated with $0-250$ $\mu$M of TMZ and data was collected at different time points [17]. The data in this picture was not used for parameter estimation. The curves were obtained by using the model with appropriate values of $\alpha$ and $K$ obtained in figure S2A.

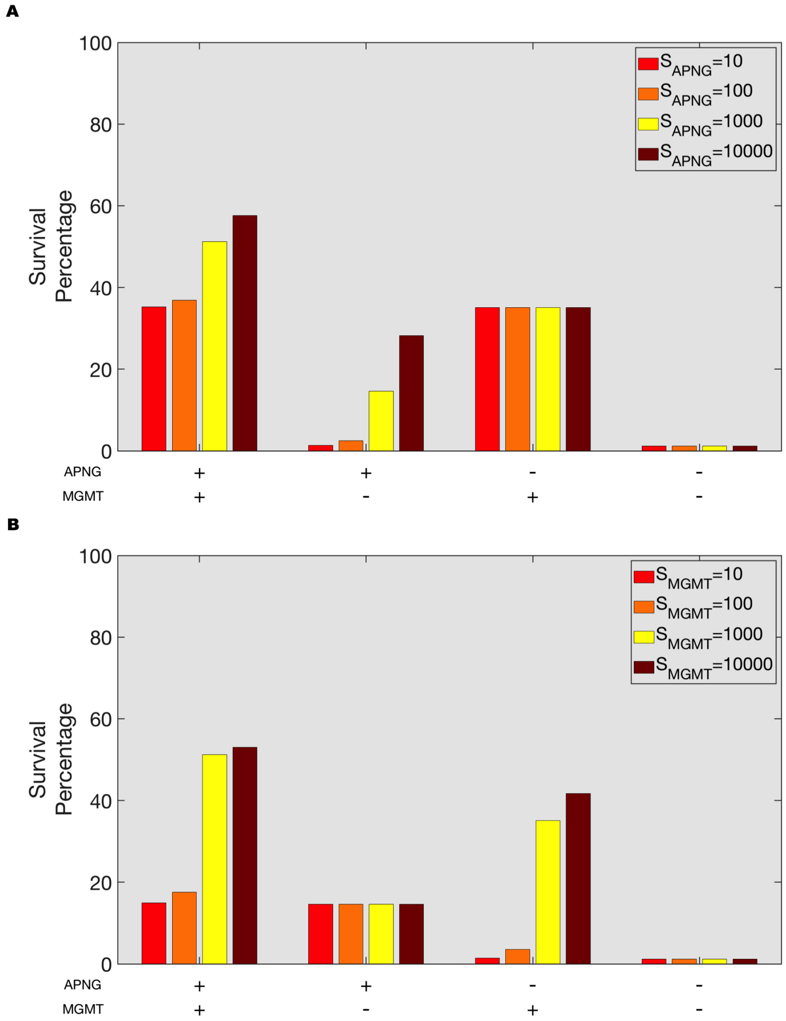


**Figure S4. Additional local sensitivity analysis.** A, Survival percentage for different enzyme expressions of APNG when APNG/MGMT are active or inhibited. B, Survival percentage for different enzyme expressions of MGMT when APNG/MGMT are active or inhibited.
